# Ali-U-Net: A Convolutional Transformer Neural Net for Multiple Sequence Alignment of DNA Sequences. A proof of concept

**DOI:** 10.1101/2025.02.26.640343

**Authors:** Petar Arsic, Christoph Mayer

## Abstract

We report a convolutional transformer neural network that is capable of aligning multiple nucleotide sequences. The neural network is based on the U-Net commonly used in image segmentation which we employ to transform unaligned sequences to aligned sequences. For alignment scenarios our Ali-U-Net neural network has been trained on, it is in most cases more accurate than programs such as MAFFT, T-Coffee, MUSCLE, and Clustal Omega, while being considerably faster than similarly accurate programs on a single CPU core. Limitations are that the neural network is still trained specifically for certain alignment problems and can perform poorly for gap distributions it has not seen before. Furthermore, the algorithm currently works with fixed-size alignment windows of 48×48 or 96×96 nucleotides. At this stage, we view our study as a proof of concept, confident that the present findings can be extended to larger alignments and more complex alignment scenarios in the near future.

## Introduction

### The Multiple Sequence Alignment problem

In molecular biology, a multiple sequence alignment is an alignment of three or more nucleotide or amino acid sequences. The rows represent different sequences and the columns position-specific residues in the sequences (Reyes et al. 2017). The aim of aligning multiple homologous sequences of e.g. nucleotides or amino acids is to maximize the number of homologous residues found in the same alignment columns of a multiple sequence alignment by adding gaps (Li 1997, Durbin et al. 1998). Homologous sequences are sequences with the same common ancestor sequence. Similarly, homologous residues are residues with a common ancestor within the homologous sequences. Homology is a hypothesis that cannot be proven, but which becomes more likely, the more similar structures, in our case the sequences, are.

Determining the best multiple or pairwise sequence alignment lies at the basis of most sequence comparisons and therefore is a key problem in bioinformatics (Smith and Waterman 1981, Li 1997, Durbin et al. 1998). With the growth of biological data over the past years, publications about multiple sequence alignment algorithms have quickly become one of the most frequently cited research areas in all of science (Van Noorden et al. 2014). This includes multiple sequence analysis for phylogenetic studies, domain analysis, and motif discovery (Durbin et al. 1998, Dotan et al. 2022). The discovery of efficient MSA algorithms is still of high interest to biologists.

The multiple sequence alignment problem belongs to the group of so-called NP-hard problems when assuming the sum-of-pairs score as the optimality criterion (Wang & Jiang 1994), with the consequence that for a larger number of sequences, heuristic methods are needed to solve the problem in a reasonable time (Elias 2006, Yang et al. 2020). This implies that a computationally adequate alignment may not always accurately reflect historical evolutionary events (Morrison 2008) and that alignment success may depend on the specific attributes of the alignment problem.

To address the multiple sequence alignment problem, various algorithms and programs have been developed to optimize alignment accuracy and efficiency (e.g. Needleman and Wunsch 1970, Hirschberger 1975, Smith and Waterman 1981, Gotoh 1982, Feng and Doolittle 1987). Common tools for aligning multiple sequences include CLUSTAL-W (Thompson et al. 1994), T-Coffee (Notredame et al. 2000), MUSCLE (Edgar 2004), and MAFFT (Katoh and Standley 2013). The most widely used heuristic is the so-called progressive alignment algorithm proposed by Feng and Doolittle (1987), in which groups of taxa are aligned in the putative reverse evolutionary order from the tips to the root, i.e. by first aligning more similar groups of sequences before aligning larger more distantly similar blocks of sequences. Ishikawa (1994), Notredame (1996), Zhang (1997), Lee et al. 2008, and Naznin et al. (2011) and many more (see references in these publications) proposed genetic algorithms for the multiple sequence alignment problem, a class of traditional machine learning algorithms.

Machine learning encompasses a variety of techniques for creating predictive models that learn patterns from data, rather than relying on explicitly programmed rules. These models can then be used to make predictions on previously unseen data. Due to advances in theoretical understanding and available hardware, artificial neural networks have become a common tool in many disciplines, including biological and medical data analyses (see e.g. Min et al. 2017, Ching et al. 2018, Greener et al. 2022, Borowiec et al. 2022, Cifci et al. 2023, Kulikov et al. 2024). A subclass of neural networks is convolutional neural networks (LeCun et al. 1989), which are frequently used for identifying features in data, such as images, and discovering motifs in DNA sequences (Yang et al. 2020).

Deep learning has been proposed for DNA sequence alignments as early as 2009 when Nielsen & Lund used a simple feed-forward network with 2 to 60 hidden neurons to align nine amino acids against a reference. Cellular neural networks (not to be confused with convolutional neural networks) have been suggested for pairwise sequence alignments by Ji et al. 2015. More recently, Jafari et al. 2019 proposed a deep reinforcement learning approach together with a long short-term memory network to determine multiple sequence alignments. Liu et al. 2023 combine reinforcement learning with a column-by-column alignment process to align longer sequences. Lall and Tallur 2023 proposed reinforcement neural networks for pairwise sequence alignments of longer sequences. Conceptually very interesting are the natural language transformer-based models proposed by Dotan et al. 2022, Dotan et al. 2024a, and Dotan et al. 2024b. They aligned up to 10 sequences and up to 1024 symbols. A disadvantage of this complex approach is its high resource consumption for longer sequences, which the authors attempt to alleviate with a tokenization approach (Dotan et al. 2024b). Their approach outperformed traditional multiple-sequence alignment programs in the investigated scenarios.

Here we suggest a novel supervised machine learning strategy for the multiple sequence alignment problem using a slightly modified U-Net (Ronneberger et al. 2015) to transform unaligned sequences to a multiple sequence alignment. The original image segmentation U-Net is built using a series of convolutional and pooling layers that encode the image (encoder branch), followed by a series of layers with upsampling operations (decoder branch), in which the resolution of the original image is reintroduced by skip connections from the encoder branch. A modified version of this architecture will be used here to transform the matrix of unaligned nucleotides into a matrix of aligned nucleotides.

The core idea behind supervised machine learning methods is to train a model by providing it with a large set of data points as input – unaligned sequences in this context – along with the expected outputs of the neural network - the aligned sequences. Due to the high amount of required training data, it will only be possible to accomplish this by using simulated data, see also Dotan et al. (2022). In the case of multiple sequence alignments, we also face the following problems. (i) For a task as complex as multiple sequence alignment, it is difficult to simulate the full variety of typical unaligned sequence datasets and (ii) for some quality scores for MSAs, the simulated alignment might not be the best possible alignment, i.e. it might be possible to find a solution that is better than the reference alignment. The latter is a general problem and it is clear that even for natural sequences an optimal pairwise identity score might favor an alignment that differs from the true alignment we would get if we were able to follow the true evolution.

## Materials and Methods

The data simulation, neural network training, and alignment prediction scripts were implemented in Python 3.10 and 3.11 using the TensorFlow library version 2.10.0 (Abadi et al., 2016).

### Encoding nucleotide sequences

In the input and output of the neural networks, the nucleotides and the gap symbol in sequences are represented by five one-hot-encoding vectors each representing one of the symbols {‘A’, ‘C’, ‘G’, ‘T’, ‘-’}. Therefore a sequence of length N is represented by a sparse matrix with the numbers {0,1} and dimensions N×5. This notation has already been used by Andreatta et al. 2016 and Almagro Armenteros et al. 2017 and is described in more detail in the Supplementary Materials and Results Section 1.1.

### Neural network architectures and implementation

Inspired by the U-Net (Ronneberger et al. 2015), a neural network architecture often used in image segmentation tasks, we devised the Ali-U-Net. We modified the original U-Net such that it can transform matrices of unaligned nucleotides to matrices of aligned nucleotides instead of transforming images to segmentation masks. Since the nucleotides are encoded by one-hot encoding vectors of size 5, the input tensor shape of the unaligned sequences was (matrix-width, matrix-height, 5) as opposed to the tensor shape (image-width, image-height, 3) in the case of an image with 3 color channels. The output shape of the Ali-U-Net was (matrix-width, matrix-height, 5) in contrast to the output shape of the U-Net which was (image-width, image-height, number-of-segmentation-classes).

The Ali-U-Net consists of five convolutional layers and four intermediate pooling layers which downsample and encode the alignment into progressively more feature maps. It then upsamples the encoded representation of sequences again through four transpose convolutions, gradually restoring the output resolution. The downsampling layers are referred to as the contracting or encoder path, with the upsampling layers being the expansion or decoder path. Through skip connections, the output of the contracting layers is concatenated to the output of expansion layers with equal dimensions (Ronneberger et al. 2015). The most important modifications of the Ali-U-Net with respect to the original U-Net are different filter kernel sizes in the convolutional layers. Whereas a kernel size of 3×3 was used in the U-Net for all convolutional layers, we obtained better results for a cascade of kernel sizes 11×11, 7×7, 5×5, 4×4, and 3×3 in the Ali-U-Net. The architecture of the main Ali-U-Net, which we also refer to as the one-branch model (B1) is illustrated in (Figure 1).

**Figure 1:**
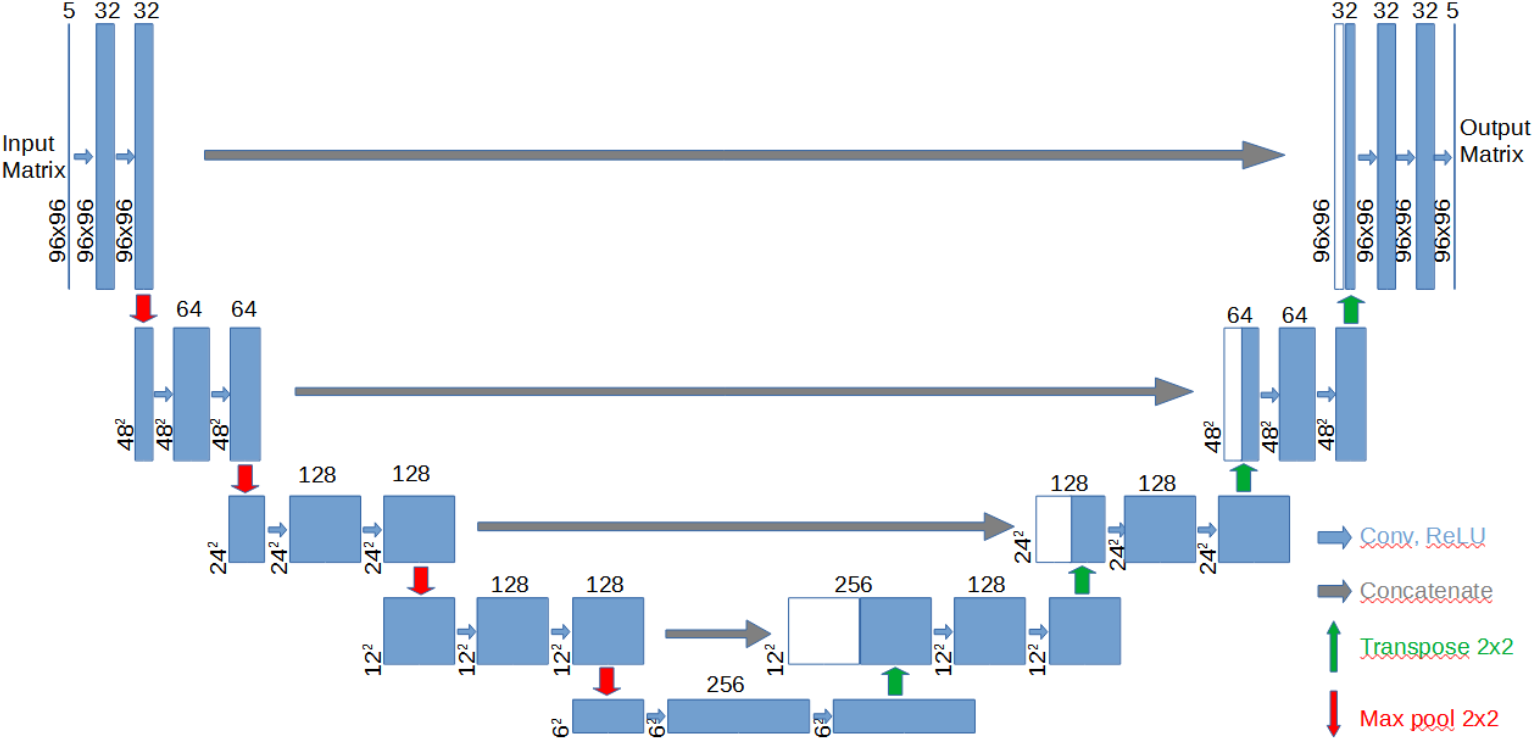
Architecture of the basic Ali-U-Net, B1 model. The color scheme and figure design were modified after (Ronneberger et al. 2015).

Besides the one-branch Ali-U-Net architecture (B1) shown in Figure 1, we also created two and three-branch variants of the Ali-U-Net, referred to as B2 and B3. In these multi-branch models, the input was fed into two, and three different U-Net branches respectively, and their outputs were concatenated before combining the final output to the five output channels for the four nucleotides and the gap symbol. The B1-, B2-, and B3-model architectures are illustrated in Supplementary Figure S5, the code is described in more detail in the Supplementary Materials and Results and is made available in the GitHub repository of the project, see the Data Availability Section.

Multiple branch neural networks can conceptually be traced back to Jacobs et al. (1991) who referred to this technique as combining the output of multiple experts. In the context of CNNs, multi-branch models were introduced to our knowledge with the introduction of the Inception-net by Szegedy et al. 2015.

The three branches used in our Ali-U-Nets were designed with the following concept in mind: The first branch with square filters extracts large-scale motifs in the alignment. The second branch with horizontal filters helps keeping the order of the nucleotides, and the third branch helps maximize the number of equal nucleotides in the alignment columns.

For all models, we chose to use “categorical-cross-entropy” as the loss function, which was also used in the original U-Net of paper by Ronneberger et al. (2015). For the activation function we used either the sigmoid (Rumelhart et al. 1986) or ReLU (Nair et al. 2010) functions for all layers, except the final layer in each branch which was always chosen to be the sigmoid activation function, see Supplementary Figure S5 and the arguments given for this choice. In some study cases, neural networks did not learn for one or both activation functions. Therefore we used activation function-specific kernel initializers to stabilize the numerical results. See Section 1.5 in the Supplementary Materials and Results for more information. We added dropout layers (Wager et al. 2013) between downsampling and between upsampling blocks. Finally, we used ADAM (Kingma & Ba 2014) as the optimizer with an initial learning rate of 0.0001.

### Generating the training dataset

Training MSA machine learning models based on the here proposed Ali-U-Net requires a large number of training datasets, i.e. pairs of unaligned and aligned sequence matrices. While first results can be achieved with a few 100,000 alignments, accurate results for specific alignment problems require about 1-10 million pairs of sequence matrices. This amount of data can only be generated using simulations.

Sequence simulations are commonly used in benchmarking workflows for multiple sequence alignment and phylogenetic reconstruction algorithms providing an *in-silico* method to test a broad range of evolutionary scenarios (Ashkenazy et al. 2017).

Simulating indels in molecular evolution is challenging and different alignment situations have very different characteristics (Cartwright 2009, Fletcher and Yang 2009). After exploring the capabilities of INDELible (Fletcher and Yang 2009) and SpartaABC (Ashkenazy et al. 2017) we decided to implement our own sequence simulator since simulated alignments were either too conserved or too difficult to align. Pairs of aligned and unaligned nucleotide sequence matrices of size 48×48 and 96×96 were generated as follows: First, backbone alignments with substitutional differences were generated. Substitutions were modeled using a profile of positional nucleotide probabilities (Reyes et al. 2017). In our simulations, we assigned to each alignment column a probability profile by randomly assigning to each of the four nucleotides one of the probabilities {90%, 4%, 3%, and 3%}. Using this column-specific profile, nucleotides were chosen randomly for all sequences. The sequences obtained in this way are related as in a star tree. Next, we added gaps to these backbone alignments. We investigated two main gap distribution patterns which were used for the 48×48 and the 96×96 alignment matrices. It should be noted that in the 48×48 and 96×96 alignment cases, three and five sequences per alignment, respectively, were selected randomly and simulated without inserting gaps. This provides a backbone for the whole alignment, avoiding gap-only columns as the potential best alignment result, and makes it easier to compare the results. The first gap distribution pattern only allows gaps at the terminal ends of the sequences and is referred to as the Ragged Sequence Ends (RSE) scenario. By this, we simulate sequences with different start and end positions. The second gap distribution pattern allows structured gap blocks within the matrix in addition to the ragged sequence ends, which we refer to as the Ragged Sequence Ends and Internal Gaps (RSE+IG) scenario.

The first two study cases we generated are called 48×48-RSE and 96×96-RSE. The terminal gaps were simulated as follows: For each sequence and both ends, a gap width was chosen randomly, in the ranges 1 to 5 for the 48×48-RSE and the range 1 to 7 in the 96×96-RSE scenario with the probabilities listed in Table 1. Additional internal gaps were added in the study cases 48×48-RSE+IG and 96×96-RSE+IG, in which the terminal gaps were simulated as before. Two internal gap blocks are inserted, one in the left and one in the right half of the alignment. For each of these blocks, a base gap width (bgw) was randomly selected from the range of possible widths from 2 to 4. In the 48×48-RSE+IG scenario, the smallest gap had a probability of 50% while the other two occurred with probabilities of 25%. In the 96×96-RSE+IG scenario, the largest gap had a probability of 50% and the smaller two had probabilities of 25% (Table 1). To increase the complexity, in each sequence, the final gap was allowed to deviate from the block structure as follows: For each sequence (except for the three/five invariant sequences), for each gap and both directions, gaps could grow by 0,1,2 bp with the probabilities 80%, 10%, 10%, respectively. Internal gaps were not allowed to be too close to the left and right border and not too close to the middle of the alignment region. Allowed ranges for these internal gaps are shown in Table 1 and the gaps are placed into this range with uniform probabilities. It should be noted that for the RSE+IG study cases, the internal gaps can fuse with the terminal gaps with a low probability. In regions simulated as gaps, nucleotides were simply overwritten by gap symbols. Unaligned sequences were generated by moving all gaps to the right end of the alignment matrix. Examples of simulated aligned and unaligned sequences are shown in the Supplementary Figures S1 to S4.

**Table 1.** Parameters used for simulating gap distribution in alignments. Listed are probabilities for terminal gap widths, internal gap block widths, and probabilities for the internal gap to vary from the block size independently for each sequence and both ends. The internal gaps are placed into the left and right half of the alignment with uniform probabilities in the specified ranges.

| Scenario | Terminal gap widths<br>RSE and RSE+IG |  | Internal gap<br>block widths<br>(RSE+IG only) |  | Internal gap<br>growth option<br>(RSE+IG only) |  | Allowed ranges for<br>Internal gaps<br>(RSE+IG only) |  |
| --- | --- | --- | --- | --- | --- | --- | --- | --- |
|  | Width | Prob. | Width | Prob. | Growth<br>options | Growth<br>prob. | Left<br>Half | Right<br>Half |
| 96x96<br>RSE,<br>RSE+IG | 0 | 12.5% | 2 | 0.25 | 0 | 0.8 | 4-46 | 50-92 |
|  | 1 | 12.5% | 3 | 0.25 | 1 | 0.1 |  |  |
|  | ... | ... | 4 | 0.50 | 2 | 0.1 |  |  |
|  | 7 | 12.5% |  |  |  |  |  |  |
| 48x48<br>RSE,<br>RSE+IG | 0 | 16.67% | 2 | 0.50 | 0 | 0.8 | 4-22 | 26-44 |
|  | 1 | 16.67% | 3 | 0.25 | 1 | 0.1 |  |  |
|  | ... | ... | 4 | 0.25 | 2 | 0.1 |  |  |
|  | 5 | 16.67% |  |  |  |  |  |  |

### Training, validation, and test datasets for the four main study cases

In this project, we first generated training, validation, and test datasets for the Ali-U-Net for the four main study cases, namely the 48×48-RSE, 48×48-RSE+IG, 96×96-RSE, 96×96-RSE+IG. For every study case, we created a training dataset with 10 million simulated alignments, a validation dataset with 1000 simulated alignments, and an independent test dataset of 1000 alignments. The validation datasets were used to determine the best model architecture and other hyperparameters. The test datasets were used for the comparison with other alignment software. The independence of the validation and test datasets is important, since in principle models and hyperparameters could be biased towards a better performance on the validation datasets.

### Training the Neural networks

The four main study cases were trained using the corresponding training dataset with 10 million and a validation dataset with 1000 pairs of nucleotide matrices. The training was conducted using a batch size of 64 and 50 epochs. During each training, we saved the best model Checkpoint, i.e. the one with the highest validation accuracy over all epochs. Furthermore, we saved the model obtained after the last training epoch. If training times are given, they were determined when using an A100 Nvidia GPU and should only be seen as a rough estimate, since training was done on shared resources.

### Evaluating Alignment Accuracy

Several optimality criteria exist for measuring and comparing the accuracy of multiple sequence alignments.

i. For the so-called sum of pairs (SoP) score there exist two alternative definitions in the literature. (ia) Most often, the SoP score is defined as the sum of all pairwise similarity scores in an alignment (Murata et al. 1985, Bacon and Anderson 1986, Wang & Jiang 1994). Depending on the data type and task, different pairwise scores could be envisaged. Normally, the sum of pairs score is determined in normalized form. (ib) Alternatively, the SoP score can be defined as the proportion of correctly aligned residues in comparison to a reference alignment, i.e. the proportion of pairs of residues found in the same alignment column in the new alignment as well as in the reference alignment (Thompson et al. 1999, Sauder et al. 2000, Edgar 2004). This score can only be used, if a correctly aligned reference is available and cannot serve as an optimality criterion for scoring alignments without a known reference. In the present study, we used the SoP score (ia) with a simple identity score for pairwise comparisons. Therefore, this score is equal to the total sum of all pairs of identical residues that occur in the same alignment columns throughout the entire alignment. This sum is normalized by dividing it by the total number of compared residue pairs, resulting in values ranging from 0 to 1.0. A value of 1.0 is obtained if all sequences are pairwise identical in regions not containing gaps.
ii. A comparatively strict score is the total percentage identity between an alignment and a reference alignment, which is assumed to be the correct alignment. In our implementation, gaps are included in the comparison. Thus, this score is 1.0 if the alignment and its reference are identical. A disadvantage of this score is that the reference alignment is assumed to be the only correct alignment, even if equally good or potentially even better alignments exist. This metric is identical to the “categorical_accuracy”, which TensorFlow determines during the training process. For simplicity, we will refer to this as the alignment accuracy.

### Comparing the Ali-U-Net with state-of-the-art alignment software

We compared the alignment results of the Ali-U-Net with four state-of-the-art alignment tools: MAFFT v7.526, T-Coffee v11.0.8, MUSCLE v5.1, and Clustal Omega v1.2.4. Default options were used for all tools except for MAFFT, where the localpair algorithm was used for the alignment problems with internal gaps, and the globalpair algorithm for those without. The Ali-U-Net and the four well-known alignment programs were used to align the 1000 sequence matrices in the test dataset for each study case. For these alignments, we determined the identity and the SoP score.

### Testing generalisability with models trained on mixtures of RSE and RSE+IG alignments

The question arose whether NNs can be trained such that one model for each matrix size performs well on both test datasets of the RSE and RSE+IG study cases. Therefore we trained Ali-U-Net_mix_ models for two additional study cases, 48×48-RSE_IG_mix_ and 96×96-RSE_IG_mix_, again using training datasets of 10 million and validation datasets of 1000 alignments. These training and validation datasets consisted of random mixtures (50%, 50%) of pairs of unaligned and aligned sequences from the RSE and RSE+IG gap distribution cases. Models trained in these additional study cases were evaluated on the test datasets of the four main study cases, so no additional test dataset was generated. The RSE_IG_mix_ study cases allow us to evaluate the next steps toward more general models.

### Performance on untrained gap distribution

In a final test, we determine the performance of the Ali-U-Net_mix_ models on a gap distribution far off what they have been trained on. For this, we simulated 1000 alignments with 1-3 unstructured gaps per alignment with a random length in the range from 1 base pair to 20% of the alignment length. These gaps were allowed to occur with equal probabilities over the full length of the sequences. We will refer to this as the unstructured gap distribution.

### Measuring the computation times for the alignment programs

Computation times were determined by aligning all 1000 unaligned matrices in the test datasets with all five alignment programs. For a fair comparison, all programs were allowed to use only a single CPU core. For the Ali-U-Net we also report the prediction times when a GPU (NVIDIA A100, CUDA version 11.7.0) was used. Computation times using a single core were determined on the same computer without sharing it with other users. The tests with GPUs were conducted on shared resources.

### Estimating hallucinations

Transformer models have a non-vanishing probability of hallucinating a wrong outcome, which in our case would be a wrong base or gap at some positions of the predicted alignment. Comparing unaligned sequences and sequences aligned with the Ali-U-Net, one can observe a low rate of wrong nucleotide bases or gaps, typically altered such that the number of equal symbols in an alignment column increases. Hallucinations in an alignment predicted with the Ali-U-Net are detected with the following method: We extract the individual sequences of the predicted alignment and remove all inserted gaps. These extracted sequences have then been aligned pairwise with the original input sequences. Pairwise alignments were determined with the Biopython module pairwise2 using the `pairwise2.align.globalms` function (Cock et al. 2009) and the alignment parameters 1, 0, -2, -1. Altered symbols can be identified as differences in this alignment. In principle and in the presence of too many hallucinations, it could be that no good alignment can be found and the number of differences is overestimated. Thus, this method provides an upper bound for the number of hallucinations in the alignment. As long as the typical number of hallucinated bases is low, which is the case for most alignments in our study cases, the estimated number of hallucinations is close to the true number of hallucinations.

## Results

### Determining the best model and activation function

First, we determined the best neural network architecture together with the best activation function. We want to note that considerable optimizations of the filter kernel sizes, the number of layers, and dropout rates have been conducted, but cannot be fully detailed here. Validation accuracy learning curves give a first indication of whether neural networks can learn efficiently during supervised training. In Figure 2 we show the validation accuracy during the training of the Ali-U-Net B3 model for the sigmoid and ReLU activation functions when using the full training datasets of 10 million simulated alignments generated for the four main study cases. Validation accuracies were determined after each epoch for the corresponding validation datasets consisting of 1000 nucleotide matrices. The validation accuracy learning curves show a good training convergence towards considerably high validation accuracies.

**Figure 2:**
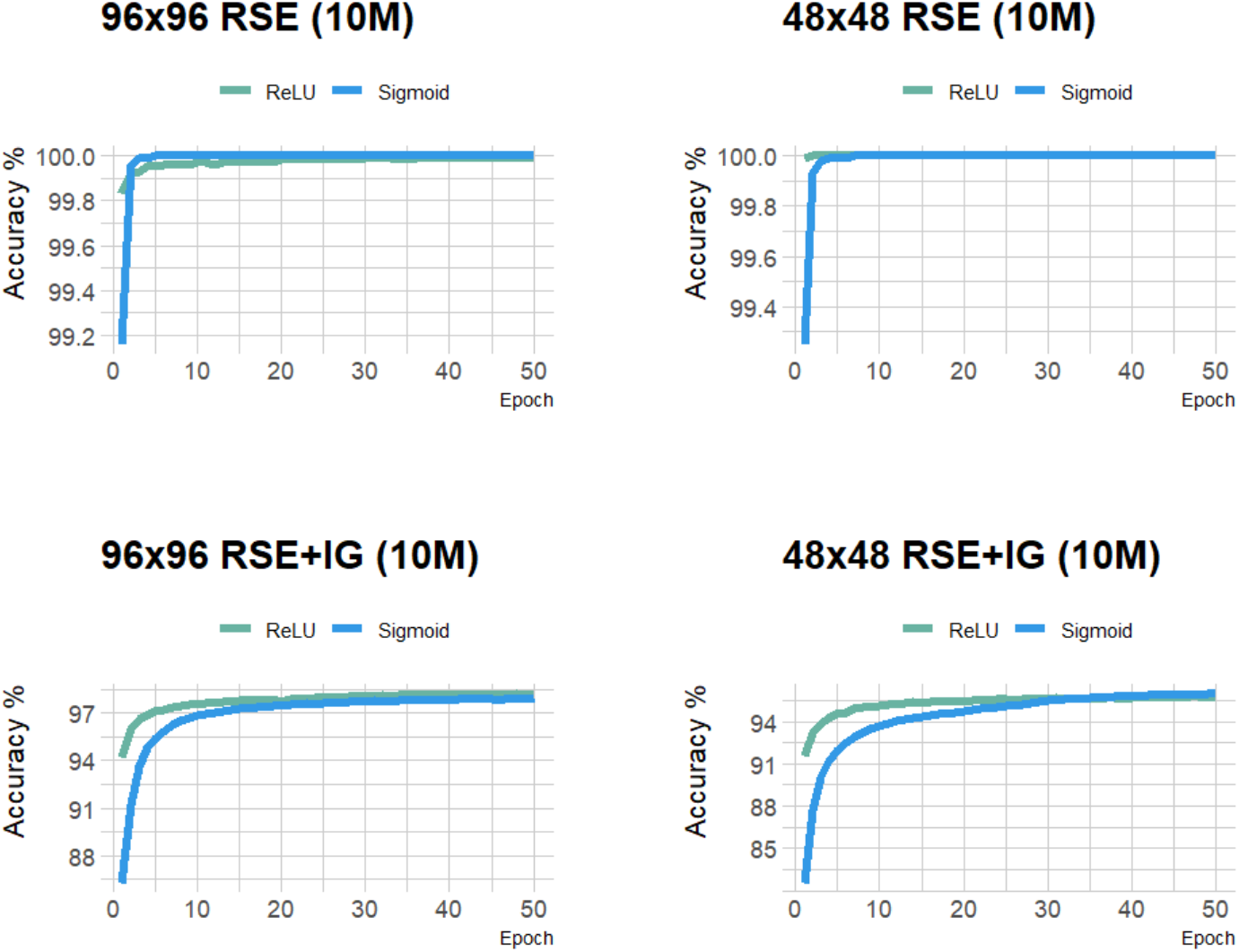
Validation accuracy learning curves for the four main study cases using the Ali-U-Net B3 model architecture. Networks were trained using the full training datasets of the four study cases and validation accuracies were determined for the corresponding validation datasets. The two lines in each subplot show the training histories under the sigmoid and ReLU activation functions.

For the 48×48 RSE scenario, an accuracy of 100% is reached after the second epoch using ReLU and the seventh epoch using the sigmoid activation. For the 96×96 RSE scenario, 100% was reached after the sixth epoch when using the sigmoid while a peak accuracy of 99.99% was attained after 30 epochs using ReLU. When internal gaps are present, the training process slows significantly. In the 48×48 RSE+IG scenario, the model achieves a peak accuracy of 95.73% with ReLU and 95.94% with sigmoid activation after 50 epochs. The 96×96 RSE+IG scenario reaches a peak accuracy of 98.17% with the ReLU and 97.93% with the sigmoid activation after 50 epochs. Altogether, the two activation functions perform similarly well. No activation function is better in all cases. ReLU is preferred due to its tendency to achieve peak results earlier (Figure 2).

Next, we compared the alignment accuracy of the B1, B2, and B3 neural network architectures for the sigmoid and ReLU activation functions. Since we achieved mean alignment accuracies of 100% for the RSE study cases after just a few epochs, we only included the two study cases with internal gaps in this comparison. Models were trained as before on the full training datasets. For the predictions, the best model was used that was found during the training. For the comparison of the models, alignments were predicted for all 1000 unaligned nucleotide matrices in the validation datasets of the study cases. For these predicted alignments, the mean and median accuracies are shown in Table 2. An example alignment predicted with the Ali-U-Net from unaligned sequences is shown in Supplementary Figures S6-S7.

**Table 2.** Median (Mdn) and mean (M) identity between predicted and reference alignments for a comparison of the model architectures B1, B2, and B3 with sigmoid and ReLU activation functions. Models were trained for 50 epochs. The “Best Epoche” column depicts the epoch in which the highest prediction accuracy was found. Median and mean accuracies were determined for the validation datasets of the two study cases. The highest results are indicated by a grey background.

| Study case<br>(RSE+IG) | Activation<br>function | Model<br>architecture | Best<br>Epoch | Training<br>time (s) | Accuracy of best model |  |
| --- | --- | --- | --- | --- | --- | --- |
|  |  |  |  |  | Mdn | M |
| 48×48 | ReLU | B1 | 46 | 122527 | 94.92 | 94.01 |
| 48×48 | sigmoid | B1 | 50 | 126560 | 94.36 | 93.50 |
| 48×48 | ReLU | B2 | 48 | 214674 | 96.48 | 95.60. |
| 48×48 | sigmoid | B2 | 49 | 213889 | 95.79 | 94.95 |
| 48×48 | ReLU | B3 | 47 | 314956 | 96.62 | 95.78 |
| 48×48 | sigmoid | B3 | 49 | 313985 | 96.79 | 95.94 |
| 96×96 | ReLU | B1 | 48 | 445579 | 97.14 | 96.85 |
| 96×96 | sigmoid | B1 | 49 | 448437 | 97.65 | 97.35 |
| 96×96 | ReLU | B2 | 50 | 693675 | 98.15 | 97.85 |
| 96×96 | sigmoid | B2 | 50 | 713078 | 98.18 | 97.90 |
| 96×96 | ReLU | B3 | 50 | 979171 | 98.45 | 98.17 |
| 96×96 | sigmoid | B3 | 50 | 993591 | 98.22 | 97.93 |

### Model for further analyses

For the 48×48 RSE+IG study case, the B3+Sigmoid architecture attains the highest mean and median accuracy whereas for the 96×96 RSE+IG study case, the B3+ReLU architecture attains the highest median and mean accuracy (Table 2).

For consistency and since NNs with ReLU train faster, all further analyses have been conducted using the B3 model architecture with ReLU as an activation function and using the model checkpoint with the highest validation accuracy during the training, i.e. the trained model obtained after the epoch listed in the “Best epoche column in Table 2.

### Comparing the Ali-U-Net with state-of-the-art alignment programs

The accuracy of alignments predicted with the Ali-U-Net was next compared to the accuracy of alignments obtained with widely used state-of-the-art alignment programs on the test datasets for the different study cases. Table 3 and Figure 3 show that the Ali-U-Net consistently attains identity scores of about 100% for the RSE study cases, which have no internal gaps. In these study cases, the Ali-U-Net clearly outperformed the alignment programs MUSCLE, T-Coffee, and Clustal Omega. It also outperforms MAFFT in the 96×96 (RSE) study case and performs equally well as MAFFT in the 48×48 (RSE) study case in which MAFFT also obtains identity scores of 100%. For the RSE+IG study cases the Ali-U-Net achieves a mean accuracy of about 96% and on average outperforms all four state-of-the-art alignment software.

**Table 3.** Comparison of the alignment accuracies of the Ali-U-Net (B3+ReLU) and four widely used alignment programs. Listed are the median (Mdn) and mean (M) identity between predicted and reference alignments for the test datasets corresponding to the four study cases.

| Algorithm | 48×48 (RSE) |  | 96×96 (RSE) |  | 48×48 (RSE+IG) |  | 96×96 (RSE+IG) |  |
| --- | --- | --- | --- | --- | --- | --- | --- | --- |
|  | Mdn | M | Mdn | M | Mdn | M | Mdn | M |
| **MAFFT** | 100.00 | 100.00 | 100.00 | 99.83 | 78.52 | 75.90 | 95.90 | 91.60 |
| **T-Coffee** | 88.46 | 85.65 | 93.64 | 88.58 | 46.92 | 54.93 | 43.26 | 51.32 |
| **Clustal Omega** | 100.00 | 86.22 | 99.07 | 86.03 | 55.53 | 51.02 | 72.31 | 67.19 |
| **MUSCLE** | 74.35 | 73.95 | 77.02 | 76.73 | 53.67 | 59.05 | 74.09 | 69.57 |
| **Ali-U-Net (B3)** | 100.00 | 100.00 | 100.00 | 99.99 | 96.48 | 95.82 | 98.05 | 96.54 |

**Figure 3:**
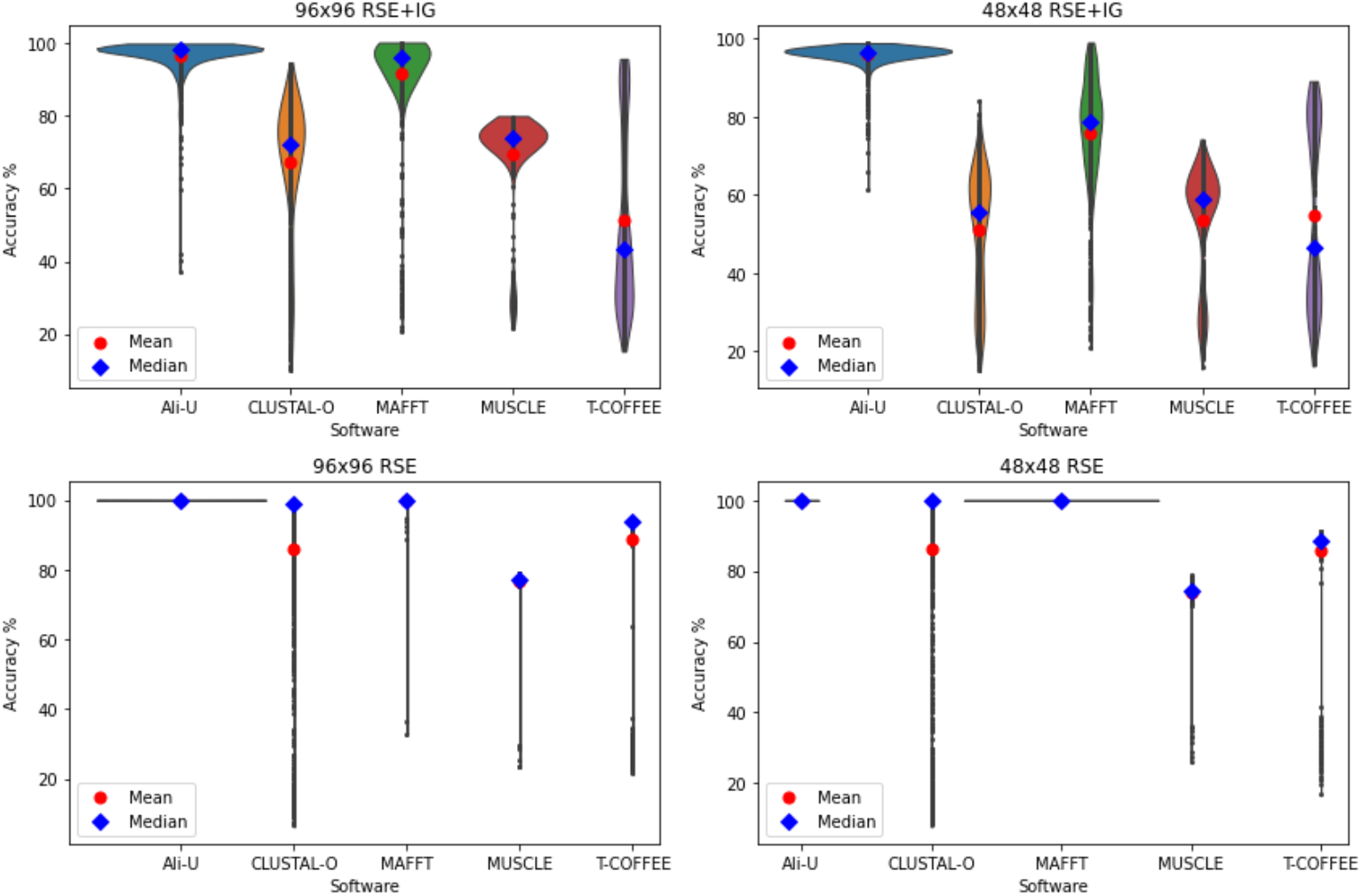
Comparison of the alignment accuracies of the Ali-U-Net (B3+ReLU) and four widely used alignment programs. The violin plots visualize the distribution of the identity scores between the predicted and reference alignments for the test datasets corresponding to the four study cases.

Besides the identity score, we also used the reference independent SoP score to compare the alignments determined with the Ali-U-Net (B3+ReLU) and the four other alignment programs. The median and mean SoP scores of the alignments in test datasets of the four main study cases are shown in Table 4 and the distributions of the SoP scores are visualized with violin plots in Figure 4. Along with the SoP scores for the inferred alignments, we also show the SoP scores for the ground truth, i.e. for the simulated reference alignments. The Ali-U-Net clearly outperforms the other alignment programs, especially in the study cases with internal gaps. Furthermore, it shows among all alignment programs mean SoP scores that are closest to the SoP score of the reference alignments.

**Table 4:** Comparison of the median (Mdn) and mean (M) SoP scores for alignments determined with the Ali-U-Net (B3+ReLU) and four widely used alignment programs on the test datasets of the corresponding study cases.

| Algorithm | 48×48 (RSE) |  | 96×96 (RSE) |  | 48×48 (RSE+IG) |  | 96×96 (RSE+IG) |  |
| --- | --- | --- | --- | --- | --- | --- | --- | --- |
|  | Mdn | M | Mdn | M | Mdn | M | Mdn | M |
| **MAFFT** | 70.00 | 69.96 | 73.41 | 73.43 | 53.02 | 53.03 | 67.29 | 67.30 |
| **T-Coffee** | 66.34 | 66.21 | 71.43 | 71.38 | 50.10 | 49.71 | 62.41 | 62.30 |
| **Clustal Omega** | 68.83 | 56.73 | 45.96 | 52.15 | 42.51 | 40.35 | 60.74 | 55.69 |
| **MUSCLE** | 69.00 | 68.93 | 73.00 | 73.02 | 51.53 | 51.76 | 66.57 | 66.63 |
| **Ali-U-Net** | 70.00 | 69.96 | 73.43 | 73.44 | 58.14 | 57.75 | 67.80 | 67.40 |
| **Simulated ref. alignment** | 70.00 | 69.96 | 73.42 | 74.44 | 57.39 | 57.23 | 68.21 | 68.27 |

**Figure 4:**
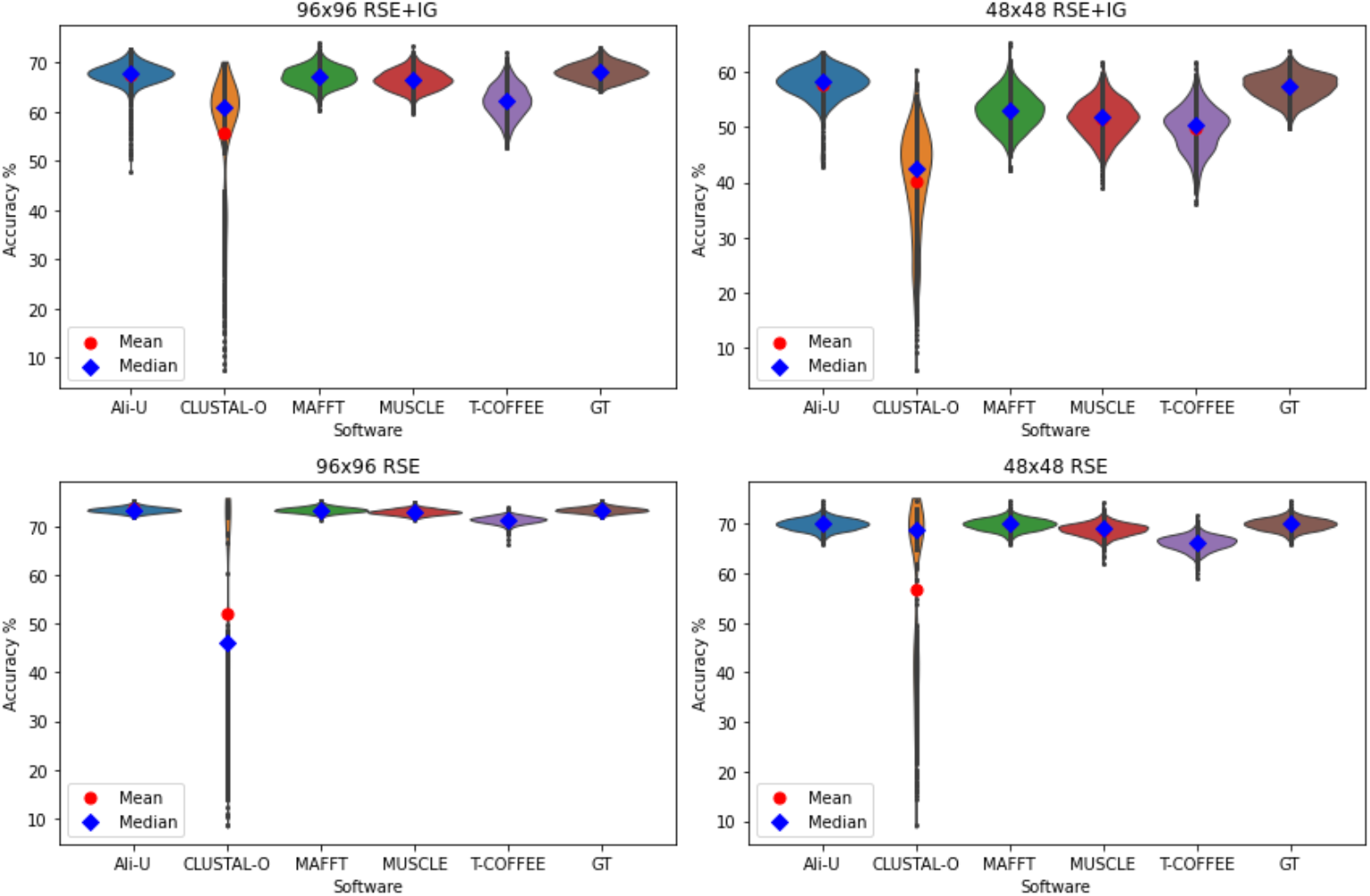
The violin plots visualize the distribution of the SoP scores for alignments determined with the Ali-U-Net (B3+ReLU) and four widely used alignment programs on the test datasets of the corresponding study cases. The ground truth depicts the distribution of the SoP scores in the simulated reference alignments.

### Performance of mixed models on the test datasets of the main study cases

To test the generalizability of the models, we evaluated the two Ali-U-Net B3_mix_ models on the four test datasets used in the RSE and RSE-IG study cases, see Table 5. In both cases, the Ali-U-Net_mix_ is only marginally less accurate than the Ali-U-Nets that have been specifically trained for the specific study cases.

**Table 5:** Comparison of the alignment accuracies of the Ali-U-Net (B3+ReLU) models trained with the study case specific training datasets (first row) and of the Ali-U-Net_mix_ models trained on mixed datasets (second row). Median and mean accuracies are evaluated for the test dataset of the study case specified in the column headers.

| Model | 48×48 (RSE) |  | 96×96 (RSE) |  | 48×48 (RSE+IG) |  | 96×96 (RSE+IG) |  |
| --- | --- | --- | --- | --- | --- | --- | --- | --- |
|  | Mdn | M | Mdn | M | Mdn | M | Mdn | M |
| **Ali-U-Net B3**<br>trained for<br>specific study<br>case | 100.00 | 100.00 | 100.00 | 99.99 | 96.48 | 95.82 | 98.05 | 96.54 |
| **Ali-U-Net B3<sub>mix</sub>**<br>trained with<br>mixed data | 100.00 | 99.99 | 99.99 | 99.98 | 96.66 | 95.96 | 97.69 | 96.40 |

### Performance of mixed models on unstructured gaps

Finally, we tested the performance of the 48×48 Ali-U-Net_mix_ when aligning sequences that were stimulated with a small number of 1-3 unstructured gaps, i.e. gaps not having a block structure. The results of this test are shown and visualized in Supplementary Materials and Results Section 3. The overall accuracy of the Ali-U-Net_mix_ drops to 95% and is only better than the MUSCLE software with 81% accuracy and worse than the other three alignment programs with about 99% accuracy. Particularly interesting is the gap character frequency distribution across the 48 alignment sites. The results show that gaps are almost exclusively added as terminal gaps since the NNs were trained only with terminal gaps and two internal blocks of gaps (Supplementary Figure S8).

### Estimated amounts of hallucinated nucleotide bases and gaps

Hallucination scores for the different study cases are shown in Table 6. Hallucination is negligible for the RSE study cases, but is clearly present in the RSE+IG scenarios with a peak value of 3.5% in the 48×48, RSE+IG study case. The lower median values compared to the mean values indicate the presence of outliers with higher amounts of hallucination. For the Ali-U-Net_mix_ models, which were also evaluated on the test datasets of the four main study cases, the hallucination is slightly higher than that of the corresponding Ali-U-Nets for the 96×96 (RSE+IG) test dataset and is clearly lower for the 48×48 (RSE+IG) test dataset.

**Table 6:** Median and mean proportion of hallucinated nucleotides or gaps for alignments predicted by the Ali-U-Net and the Ali-U-Net_mix_ models. For this test, we used the four Ali-U-Net B3 models trained for the four study cases and evaluated their predictions on the test datasets of the corresponding study cases. For the Ali-U-Net_mix_ B3 models, there exists only one variant for 48×48 and one for 96×96 input matrices. These NNs were evaluated on the four test datasets specified in the table columns.

| Model | 48×48 (RSE) [%] |  | 96×96 (RSE) [%] |  | 48×48 (RSE+IG) [%] |  | 96×96 (RSE+IG) [%] |  |
| --- | --- | --- | --- | --- | --- | --- | --- | --- |
|  | Mdn | M | Mdn | M | Mdn | M | Mdn | M |
| **Hallucination for Ali-U-Net** | 0.00 | 0.00 | 0.00 | 0.00 | 3.08 | 3.52 | 1.53 | 2.15 |
| **Hallucination for Ali-U-Net<sub>mix</sub>** | 0.00 | 0.00 | 0.01 | 0.01 | 2.77 | 3.18 | 1.66 | 2.39 |

### Computation times of the Ali-U-Net B3 model and other alignment programs

The computation times for determining 1000 alignments using the Ali-U-Net and four other alignment programs are shown in Table 7 and are visualized in Figure 5. The fastest alignment program is Clustal Omega followed by the Ali-U-Net, which is consistently the second fastest program on a single CPU core. Using a GPU, the Ali-U-Net outperforms Clustal Omega in the 96×96 study cases. For the Ali-U-Net, computing times increase by a much smaller factor than the other programs when increasing the matrix size from 48×48 to 96×96. When using a GPU, the differences between the matrix sizes are minimal.

**Table 7:** Comparison of computing times needed to determine 1000 alignments with the Ali-U-Net and other alignment programs on a single computing core, if not stated otherwise. The exponents are only a rough estimation, since for accurate results more comparisons would be required.

| Algorithm | 48×48<br>(RSE)<br>[s] | 96×96<br>(RSE)<br>[s] | Exponent<br>x in<br>$t_1=t_0 \cdot 2^x$ | 48×48<br>(RSE+IG)<br>[s] | 96×96<br>(RSE+IG)<br>[s] | Exponent<br>x for<br>$t_1=t_0 \cdot 2^x$ |
| --- | --- | --- | --- | --- | --- | --- |
| MAFFT | 362 | 723 | 1.0 | 336 | 674 | 1.0 |
| T-Coffee | 2062 | 25106 | 3.6 | 2151 | 22346 | 3.4 |
| Clustal Omega | 53 | 251 | 2.2 | 44 | 237 | 2.4 |
| MUSCLE | 611 | 6633 | 3.4 | 472 | 6293 | 3.7 |
| Ali-U-Net (B3)<br>CPU 1core | 207 | 335 | 0.7 | 238 | 340 | 0.5 |
| Ali-U-Net (B3)<br>GPU | 66 | 67 | 0.02 | 65 | 67 | 0.04 |

**Figure 5:**
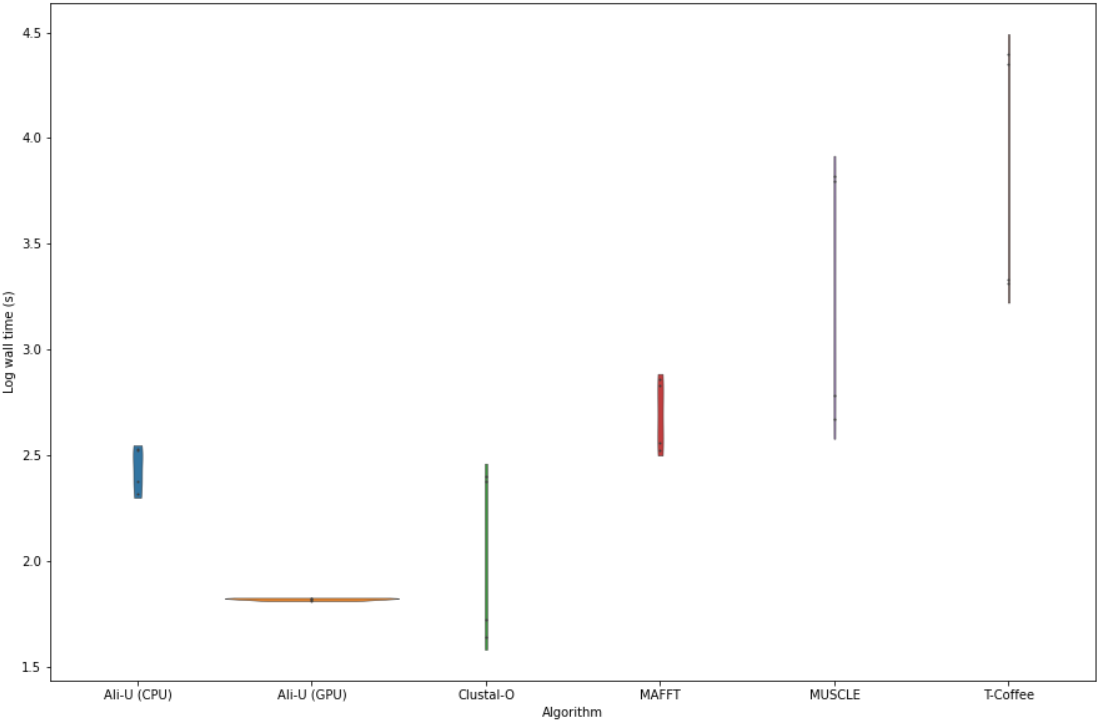
Violin plots for computing times for 1000 alignments for the Ali-U-Net and other alignment software on a single computing core.

## Discussion

We provide a proof of concept that transformer-based neural network architectures of the U-Net type can be a very accurate and efficient method for conducting multiple sequence alignments. We showed that for trained gap distribution scenarios, the Ali-U-Net can outperform traditional multiple sequence alignment programs such as MAFFT, MUSCLE, Clustal Omega, and T-Coffee in terms of total alignment accuracy and SoP score. The Ali-U-Net achieves a mean accuracy of 100% on gap distributions with only terminal gaps (Table 3). Among the four traditional alignment programs only MAFFT achieved this at the cost of 115% and 75% longer mean computing times for the 96×96 (RSE) and 48×48 (RSE) study cases (Table 7), respectively. In terms of computational speed, the Ali-U-Net was only outperformed by Clustal Omega which in all scenarios shows a much lower mean accuracy than the Ali-U-Net and MAFFT. These results indicate that the Ali-U-Net should be ideally suited for efficiently aligning a sequence to a reference as in the case of sequence mapping problems. On trained scenarios with internal gaps, the Ali-U-Net still outperforms the other programs in terms of accuracy. It is again faster than the other programs except for Clustal Omega, which also for internal gaps was the least accurate alignment program.

### The Ali-U-Net method still has strong limitations

i. It is limited to fixed-size nucleotide matrices of 48×48 or 96×96. Also other machine learning approaches for the multiple sequence alignment problems have strong limitations for the number of sequences and sequence lengths (Ji et al. 2015, Jafari et al. 2019, Lall & Tallur 2023, Dotan et al. 2024b). Dotan et al. 2024b proposed a tokenizer and k-mer approach for handling longer sequences. We believe that in the future multi-step reasoning approaches currently tested in large language models will be the best solution for piecing together very long alignments.
ii. All transformer and generator models show the tendency to hallucinate results. In our case, this means that the Ali-U-Net sometimes changes nucleotides and gaps - in most cases with the result of increasing the SoP score. Hallucination will be discussed in more detail below.
iii. Currently, and in order to be able to outperform the other multiple sequence alignment programs, the Ali-U-Net has to be trained for particular gap distributions, a restriction that also affects other machine learning methods (Dotan et al. 2022). Deviations from the gap distributions used during training lead to lower accuracies and higher rates of hallucinations, see Supplementary Materials and Results Section 2.2). This will also be discussed in more detail below.

### Best Neural Network Architecture

In this project, we optimized three neural network architectures by adjusting the number of convolutional layers, number of filters, and filter kernel sizes, details of which are not provided here. Then we compared the three architectures of the Ali-U-Net (B1, B2, B3) with one, two, and three internal branches, using either sigmoid or ReLU as activation function. We found that the architecture has a much higher influence on the alignment accuracy than the activation function (Table 2). Increasing the number of branches from one to three led to a clear and steady increase in alignment accuracies. This suggests that there is room for further improvements of the Ali-U-Net by further increasing the complexity of the model.

Our results show that the activation function only has a small influence on the final accuracy, with the ReLU activation function achieving marginally better final accuracies than the sigmoid activation function in most but not all scenarios. Due to the small differences, we cannot consider these to be significant with the tests we have conducted. A noticeable advantage of ReLU over sigmoid is faster training. The Ali-U-Net variants using ReLU reach the optimum with fewer training epochs than when using the sigmoid activation function. This result was found already for the first deep neural networks such as the Alex-Net (Krizhevsky et al. 2017). Interestingly, using the sigmoid activation in the last layer of each branch (see Supplementary Materials and Results Section 1.4.1) led to higher accuracies even in the NNs for which we used ReLU consistently for all other layers. Detailed results are not shown. Generally, using different activation functions is uncommon. It could be that using interspersed sigmoid activations is an option for optimizing NNs that is often overlooked.

### Comparing Computation Times

With growing amounts of available data, computing times are an important factor in biological sequence analysis. Different alignment algorithms have different time complexities typically stated as O(N^x^L^y^), where N is the number of sequences, L is the length of the alignment, and x and y are positive exponents. The term in parentheses describes asymptotic run times for large N and L. For short alignments, the run time can scale differently. The progressive alignment algorithm used in MAFFT has a time complexity of O(N^2^L) + O(NL^2^) (Katoh & Toh 2008), and the one used in MUSCLE of O(N^3^) + O(NL^2^) (Edgar 2004). The total runtime of Clustal Omega has a time complexity of O(N^2^) (Katoh et al. 2002) and T-Coffee of O(N^3^) (Notredame et al. 2000). The exponents we find for the different programs differ from their expected values (Table 7), most probably since the time complexity describes the asymptotic run time and the dimensions of our nucleotide matrices are small so that contributions from smaller exponents might dominate. For the range from N=48 to N=96, the Ali-U-Net has an empirical time complexity of almost O(√N) on a single core and constant time complexity when using the GPU (Table 7). In future projects, we will explore the utility of the Ali-U-Net on larger nucleotide and amino acid matrices. Altogether we find that training times of the Ali-U-Net are high and require a GPU, while prediction times are highly competitive and in most cases shorter than for other alignment software even when using only a single CPU core. For the end-user, only prediction times are relevant, if trained models are available. For this project, trained models can be downloaded from Zenodo, see Data Availability for details.

### Hallucinations in predicted alignments

Hallucination is a major concern in generative artificial intelligence (see e.g. Ji et al. 2023). Also in this project, we find a considerable tendency of the neural networks to modify the data in the direction it was trained. At this point, it should be emphasized that the SoP score can improve due to hallucinated nucleotides or gaps, but the total accuracy of the alignment gets worse in the presence of hallucinations. This excludes the possibility that the Ali-U-Net outperforms the other methods in terms of the total accuracy by modifying the sequences. The reason the Ali-U-Net alignments have a slightly higher SoP than the simulated alignments in the 48×48 (RSE+IG) study case might be due to the effect of hallucinations. Not surprisingly, the amount of hallucination increases in most cases with the complexity of the problem. While the proportion of wrong bases or gaps is <0.01% in the study cases with sequences with only terminal gaps, it increases to about 3.5% in the 48×48 (RSE+IG) study case. It is interesting and seems counterintuitive why this number is lower in the 96×96 (RSE+IG) study case. The 96×96 (RSE+IG) study case might, despite its greater sequence count and length, be easier for the net to align than its 48×48 counterpart, due to a smaller proportion of the alignment being gaps.

If the two Ali-U-Net_mix_ models, trained on RSE and RSE+IG training alignments, are evaluated on the test datasets of the four main study cases, this leads to a decreased amount of hallucination in the 48×48 RSE+IG case and a slightly increased amount in the 96×96 RSE+IG case (Table 6). Alignment accuracies with the Ali-U-Net_mix_ models are almost as good as that of the Ali-U-Net models for the plain study cases, indicating that a single model can be trained to conduct alignments on very different scenarios.

### Effects of training data being very different from test data

We found that the Ali-U-Net_mix_ models did not perform well in aligning what we called unstructured gaps (Supplementary Table S1). This gap distribution differed strongly from the distributions the two Ali-U-Net_mix_ models have been trained on. Interestingly, this scenario led to shifting the few unstructured gaps to the terminal ends of the sequences, where the training data contains a lot of gaps, see Supplementary Materials and Results Section 2.2. Looking at individual sequences, we found that in sequences in which gaps were moved to the terminal ends, a much larger number of hallucinations were detected (Supplementary Figures S8-S9). We expect that training a model on unstructured gaps will restore the strong performance of the Ali-U-Net in relation to the other alignment software. Altogether, we found that alignment neural networks need to be trained on all sequence characteristics that may occur in the data they are intended to make predictions on. The underlying problem is not new. Katoh et al. (2002) wrote that an “appropriate alignment algorithm depends on the nature of the sequences to be aligned”. Dotan et al. (2022) state that “BetaAlign heavily depends on the simulator” and argue that the use of machine learning methods in multiple sequence alignment might shift the customization pressure from the aligner to the simulator. Very generally this means that alignment simulators have to be developed that reflect the full variety of naturally occurring sequence characteristics as closely as possible. Then, either many models need to be trained specifically for different alignment problems, or preferably, fewer models on combinations of alignment problems if possible.

### Putative Functions of the Encoder and Decoder Paths in the Ali-U-Net

The main idea behind the Ali-U-Net is to transform unaligned sequences into aligned sequences similar to transforming images into segmented images. For the input alignments, the encoder path finds the common sequence patterns and encodes this information in the feature maps. The decoder path aligns the structures the feature maps contain and puts the full sequence information back together, ideally at the right positions in the alignment. Here we see an important difference between the segmentation and the alignment problem. In the case of alignments, patterns are moved and it was not a priori clear that the Ali-U-Net would be able to accomplish this. The ability to align similar regions might vanish with increasing distances in the input sequences, but this needs to be studied in more detail. In the end, we expect to see two different, but connected limits, (i) a sequence length limit and (ii) a coherence identification limit which restricts the possibility to identify similarities over large distances and to insert long gaps. Both limits are influenced by the neural network architecture and require further investigation in future work.

### Outlook and Potential for Improvement

Recently there has been significant progress in developing one-shot segmentation neural networks such as the Segment Anything model (Kirillov et al. 2023). Drawing further from the parallelism between segmentation and multiple sequence alignment, a next step could be to test the utility of the model architecture used in the Segment Anything model for training one-shot multiple sequence alignment neural networks. Since the most complex models we have investigated are those predicting the best alignments, we see potential for future improvements in terms of neural network architecture and hyperparameter optimization. The two main goals that need to be addressed are making the model more general and producing fewer hallucinated bases.

Beyond the proposed application, the presented approach could inspire researchers to extend the utility of U-Net-type neural networks beyond image processing.

## Conclusion

On gap distribution the Ali-U-Net has been trained on, it can yield alignments that are as accurate or more accurate than those obtained with existing multiple sequence alignment programs such as MAFFT, MUSCLE, T-Coffee, and Clustal Omega. Furthermore, the Ali-U-Net is faster than all programs except Clustal Omega, which is the least accurate among the tested programs. For the Ali-U-Net, three main limitations exist: (i) At the moment, the nucleotide matrices have a fixed size. (ii) The Ali-U-Net only performs well on gap distributions it has been trained on. (iii) Especially for gap distributions it has not been trained on, the Ali-U-Net tends to modify nucleotide bases and gaps, a well known tendency called hallucination that has also been observed in other applications of neural networks. We see the current project as a proof of concept, which still needs further development. In particular, we expect that even better transformer neural network architectures might exist for the multiple sequence alignment problem. Furthermore, we are convinced that aligning sequences to a reference and thereby filtering and classifying sequences can readily be accomplished with the ideas provided in this work.

## Supporting information

Supplementary Materials and Results

## Data Availability

The code for all scripts used in the present project is available from the GitHub repository https://github.com/parsec3/Ali-U-Net in release 1. The code and a more detailed explanation of the neural networks are available in the Supplementary Materials and Results. Trained neural networks are available from the Zenodo repository https://zenodo.org/records/14876519.

## Acknowledgments

The authors gratefully acknowledge the access to the Marvin cluster of the University of Bonn without which the current project would not have been possible. We are also thankful for support provided by the HPC@HRZ Team of the University of Bonn. Some preliminary computations and final evaluations were conducted using the high-performance computing resources of the Leibniz Institute for the Analysis of Biodiversity Change, Bonn.

