## Supplementary Materials and Results for "Ali-U-Net: A Convolutional Transformer Neural Net for Multiple Sequence Alignment of DNA Sequences. A proof of concept"

#### 1 Supplementary Materials

The data simulation, training, and prediction scripts as well as the neural networks (NNs) were implemented in Python 3.10 and 3.11 using the TensorFlow (Abadi et al., 2016) and Keras (Chollet et al., 2015) libraries together with the Numpy library (Harris et al., 2020) for handling the large number of training alignments.

##### 1.1 Encoding the Sequences

Nucleotides and the gap symbols in sequences are represented by five orthogonal one-hot-encoding vectors, each representing one of the symbols {'A', 'C', 'G', 'T', '-'}.

More specifically, A is represented by [1,0,0,0,0], C by [0,1,0,0,0], G by [0,0,1,0,0], T by [0,0,0,1,0], and a gap ("-") by [0,0,0,0,1]. This notation has already been used by Andreatta et al. 2016 and Almagro Armenteros et al. 2017.

Example: The sequence "GT-AAC" is encoded as follows:

```
[
   [0, 0, 1, 0, 0], # G
   [0, 0, 0, 1, 0], # T
   [0, 0, 0, 0, 1], # -
   [1, 0, 0, 0, 0], # A
   [1, 0, 0, 0, 0], # A
   [0, 1, 0, 0, 0]  # C
]
```

#### 1.2 Example alignments generated with our alignment simulator

The algorithm used to simulate alignments is explained in detail in the main text. To get an idea of how alignments generated in this way look, we show examples of pairs of simulated aligned and the corresponding unaligned sequence in Figures S1-S4.

#### 1.3 Generating and storing the training data

The training data was created with a separate script and stored on the hard drive. Since training dataset files were up to 170 Giga Byte (GB) large and stored in an encoded format for reducing the file size, handling the data was challenging. Even on compute nodes with a large amount of RAM and Nvidia GPUs with 80GB of RAM, the data could only be handled by loading it incrementally from TensorFlow record files and supplying it to the TensorFlow dataset using a generator. For details see the AlignmentSim*.py scripts in the GitHub repository https://github.com/parsec3/Ali-U-Net of this project.

#### 1.4 Implementation and design of the three Ali-U-nets.

The most important characteristics of the Ali-U-Nets are already explained in the main text. The main Ali-U-Net model architecture was inspired by the U-Net of Ronneberger et al. 2015. We have adapted the U-Net so that it can transform matrices of unaligned nucleotide sequences to matrices of aligned nucleotide sequences, instead of images with different color channels to matrices containing segmentation class information. During the process of determining the optimal NN architectures, we tested a wide range of numbers of layers, kernel sizes and numbers of kernels and other hyperparameters to optimize the alignment success of the three model architectures. Further improvements are expected to be possible, since (i) aligning nucleotide sequences is a complex task and (ii) the training process is computationally demanding and only a limited number of hyperparameter combinations could be tested.

The full code for training and prediction scripts is available in the GitHub repository https://github.com/parsec3/Ali-U-Net. Trained neural networks can be found in HERE-URL. Users can train their own neural networks using the training scripts provided in the GitHub repository.

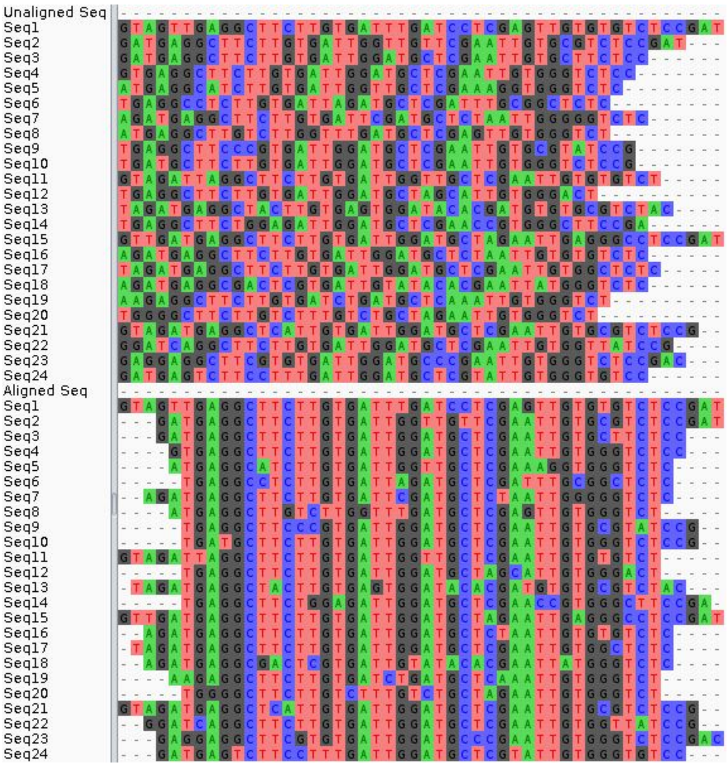


**Figure S1**: Example showing the first 24 sequences of a randomly selected pair of unaligned and aligned sequences from the 48x48 RSE training dataset.

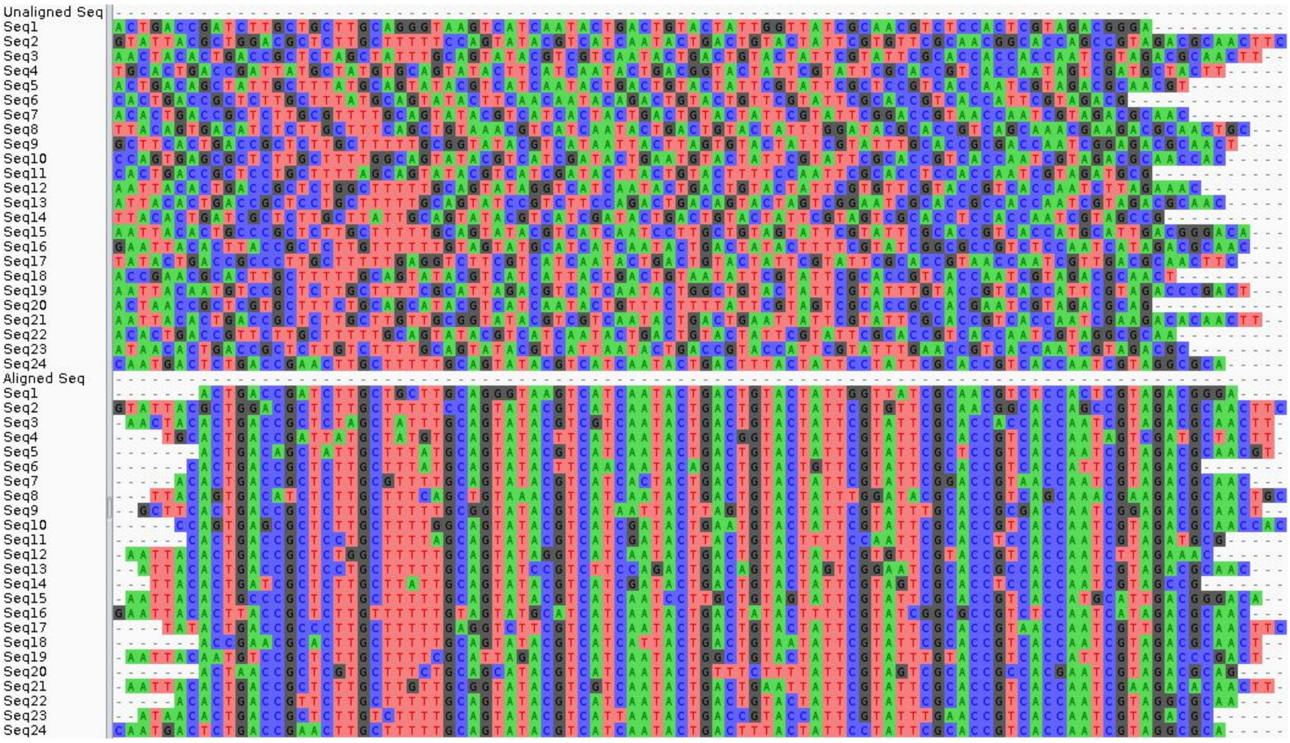


**Figure S2:** Example showing the first 24 sequences of a randomly selected pair of unaligned and aligned sequences from the 96x96 RSE training dataset.

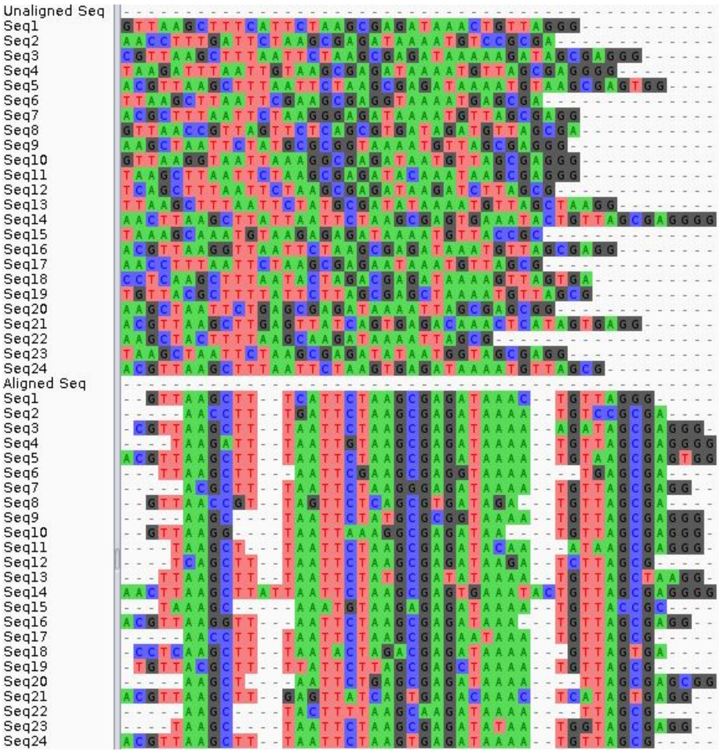


**Figure S3**: Example showing the first 24 sequences of a randomly selected pair of unaligned and aligned sequences from the 48x48 RSE+IG training dataset.

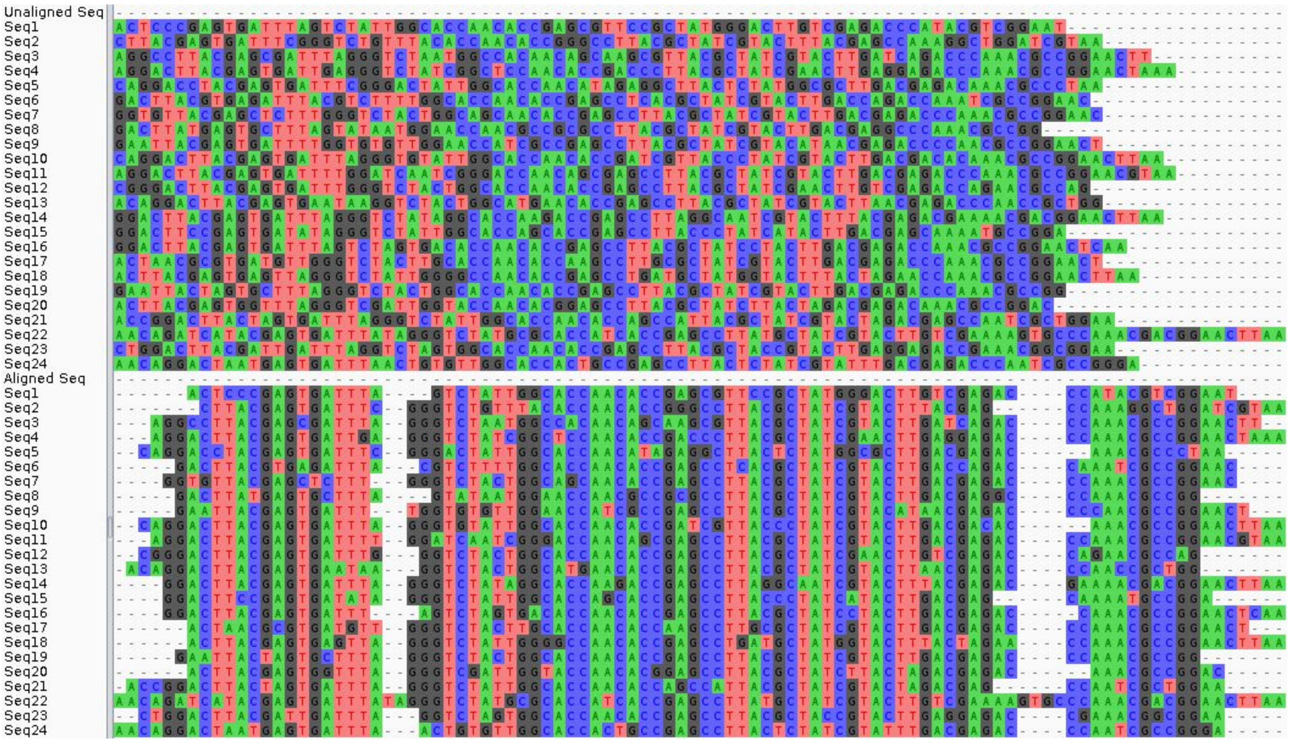


**Figure S4**: Example showing the first 24 sequences of a randomly selected pair of unaligned and aligned sequences from the 96x96 RSE+IG training dataset.

##### 1.4.1 The One-Branch Ali-U-Net

The architecture of the one-branch Ali-U-Net (B1) is very similar to the original U-Net in the paper by Ronneberger et al. (2015). The main differences are the following: (i) Filter kernel sizes of the convolutional layers are chosen to be larger in the Ali-U-Net than in the U-Net. While Ronneberger et al. (2015) used a kernel size of 3×3 for all convolutional layers, we found that larger kernel sizes increased the accuracy. Therefore kernel sizes of 11×11, 7×7, 5×5, 4×4, and 3×3 were used in convolutional layers in the encoder path. During upsampling, using `Conv2DTranspose` layers, the kernel sizes increased in reverse order in the four upsampling steps, namely 4×4, 5×5, 7×7, and 11×11. Note, that the shapes in the max-pooling layer as well as in the transpose convolutional layers remained (2,2). (ii) In the present project we used `padding="same"` as an option for all convolutional and transpose convolutional layers instead of `padding="valid"` as in the original U-Net. (iii) We use two Conv2D layers with kernel size (1, 1) and 5 kernel filters at the end of the NN. The first one uses sigmoid as an activation function, even in the cases where ReLU was chosen for all other activations. Normally, it is recommended to use the same activation function in all layers. We introduced this layer by accident. When using ReLU instead of sigmoid in this layer, the accuracy decreased noticeably by about 0.3 percent points in the 96x96-IG study case. We also tested removing the layer entirely, which resulted in a decreased accuracy. Therefore we kept the additional layer. As a consequence, in the NNs where we say that we used ReLU, we used ReLU except in this single layer. In the NNs where we used sigmoid, we used sigmoid in all layers. The model architecture of the B1 model is illustrated in Fig. 1 as well as in Fig. S5 (B1). The following code of the B1 model was used in the 48×48 as well as the 96×96 study cases. Output shapes in the following code depend on whether the input size is 48×48 or 96×96. To understand the encoder and decoder path, we added the output shape of some layers as comments for the 48×48 study cases.

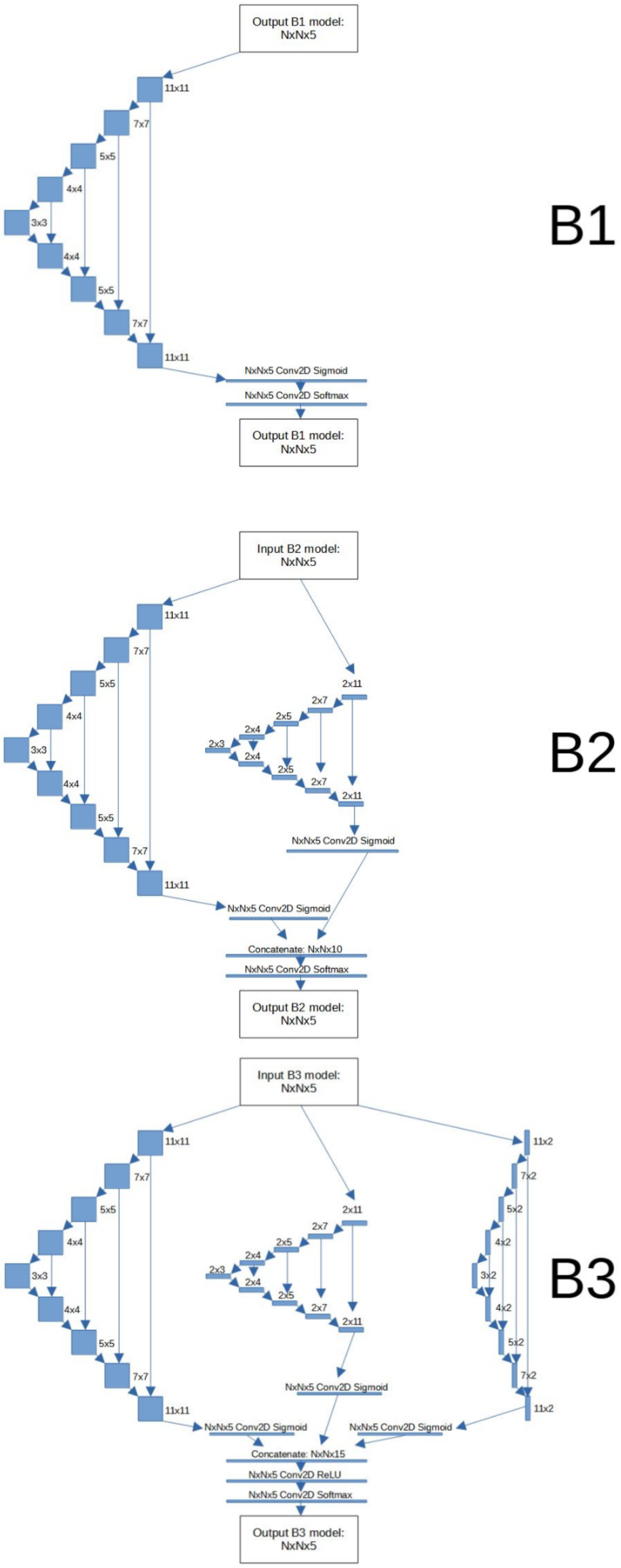


**Figure S5:** *Visualization of the three Ali-U-Net model architectures. B1: One-branch model, B2: Two-branch model, B3: Three-branch model.* The same input alignment is provided to either one, two, or three branches. The output of multiple branches is concatenated and reduced such that it has the same shape as the input matrix with five channels. For details see Section 1.1 Encoding the Sequences. Unless indicated otherwise, the same main activation function (sigmoid or ReLU) was used for all layers.

##### Code of the B1 Model Architecture

```
import tensorflow as tf
numRows = 48       # OR 96
numCols = 48       # OR 96
act_fun = “sigmoid” # OR “relu”
act_init = “glorot_normal” # OR “he_normal”

inputs = tf.keras.layers.Input(shape=(numRows, numCols, 5))
# Contraction path
c1 = tf.keras.layers.Conv2D(32, (11, 11), activation=act_fun,
kernel_initializer=act_init, padding='same')(inputs)
# Output shape: 48x48x32
c1 = tf.keras.layers.Dropout(0.1)(c1)
c1 = tf.keras.layers.Conv2D(32, (11, 11), activation=act_fun,
kernel_initializer=act_init, padding='same')(c1)
# Output shape: 48x48x32
p1 = tf.keras.layers.MaxPooling2D((2, 2))(c1)
# Output shape: 24x24x32

c2 = tf.keras.layers.Conv2D(64, (7, 7), activation=act_fun,
kernel_initializer=act_init, padding='same')(p1)
# Output shape: 24x24x64
c2 = tf.keras.layers.Dropout(0.1)(c2)
c2 = tf.keras.layers.Conv2D(64, (7, 7), activation=act_fun,
kernel_initializer=act_init, padding='same')(c2)
# Output shape: 24x24x64
p2 = tf.keras.layers.MaxPooling2D((2, 2))(c2)
# Output shape: 12x12x64

c3 = tf.keras.layers.Conv2D(128, (5, 5), activation=act_fun,
kernel_initializer=act_init, padding='same')(p2)
# Output shape: 12x12x128
c3 = tf.keras.layers.Dropout(0.2)(c3)
c3 = tf.keras.layers.Conv2D(128, (5, 5), activation=act_fun,
kernel_initializer=act_init, padding='same')(c3)
# Output shape: 12x12x128
p3 = tf.keras.layers.MaxPooling2D((2, 2))(c3)
# Output shape: 6x6x128

c4 = tf.keras.layers.Conv2D(128, (4, 4), activation=act_fun,
kernel_initializer=act_init, padding='same')(p3)
# Output shape: 6x6x128
c4 = tf.keras.layers.Dropout(0.2)(c4)
c4 = tf.keras.layers.Conv2D(128, (4, 4), activation=act_fun,
kernel_initializer=act_init, padding='same')(c4)
# Output shape: 6x6x128
p4 = tf.keras.layers.MaxPooling2D(pool_size=(2, 2))(c4)
# Output shape: 3x3x128

c5 = tf.keras.layers.Conv2D(256, (3, 3), activation=act_fun,
kernel_initializer=act_init, padding='same')(p4)
# Output shape: 3x3x256
c5 = tf.keras.layers.Dropout(0.3)(c5)
c5 = tf.keras.layers.Conv2D(256, (3, 3), activation=act_fun,
kernel_initializer=act_init, padding='same')(c5)
#Output shape: 3x3x256

#Expansive path
u6 = tf.keras.layers.Conv2DTranspose(128, (2, 2), strides=(2, 2),
```

```
padding='same')(c5)
# Output shape: 6x6x128
u6 = tf.keras.layers.concatenate([u6, c4])
# Output shape: 6x6x256
c6 = tf.keras.layers.Conv2D(128, (4, 4), activation=act_fun,
kernel_initializer=act_init, padding='same')(u6)
# Output shape: 6x6x128
c6 = tf.keras.layers.Dropout(0.2)(c6)
c6 = tf.keras.layers.Conv2D(128, (4, 4), activation=act_fun,
kernel_initializer=act_init, padding='same')(c6)
# Output shape: 6x6x128

u7 = tf.keras.layers.Conv2DTranspose(128, (2, 2), strides=(2, 2),
padding='same')(c6)
# Output shape: 12x12x128
u7 = tf.keras.layers.concatenate([u7, c3])
# Output shape: 12x12x256
c7 = tf.keras.layers.Conv2D(128, (5, 5), activation=act_fun,
kernel_initializer=act_init, padding='same')(u7)
# Output shape: 12x12x256
c7 = tf.keras.layers.Dropout(0.2)(c7)
c7 = tf.keras.layers.Conv2D(128, (5, 5), activation=act_fun,
kernel_initializer=act_init, padding='same')(c7)
# Output shape: 12x12x128

u8 = tf.keras.layers.Conv2DTranspose(64, (2, 2), strides=(2, 2),
padding='same')(c7)
# Output shape: 24x24x64
u8 = tf.keras.layers.concatenate([u8, c2])
# Output shape: 24x24x128
c8 = tf.keras.layers.Conv2D(64, (7, 7), activation=act_fun,
kernel_initializer=act_init, padding='same')(u8)
# Output shape: 24x24x128
c8 = tf.keras.layers.Dropout(0.1)(c8)
c8 = tf.keras.layers.Conv2D(64, (7, 7), activation=act_fun,
kernel_initializer=act_init, padding='same')(c8)
# Output shape: 24x24x64

u9 = tf.keras.layers.Conv2DTranspose(32, (2, 2), strides=(2, 2),
padding='same')(c8)
# Output shape: 48x48x32
u9 = tf.keras.layers.concatenate([u9, c1], axis=3)
# Output shape: 48x48x64
c9 = tf.keras.layers.Conv2D(32, (11, 11), activation=act_fun,
kernel_initializer=act_init, padding='same')(u9)
c9 = tf.keras.layers.Dropout(0.1)(c9)
c9 = tf.keras.layers.Conv2D(32, (11, 11), activation=act_fun,
kernel_initializer=act_init, padding='same')(c9)
# Output shape: 48x48x32

outputs = tf.keras.layers.Conv2D(5, (1, 1), activation='sigmoid')(c9)
# Output shape: 48x48x5
a = tf.keras.layers.Conv2D(5, (1, 1), activation="softmax")(outputs)
B1_model = tf.keras.Model(inputs=[inputs],outputs=a)
# Output shape: 48x48x5

# Create and compile the model# Adam is chosen as an optimizer with a
learning rate of 0.0001
```

```
opt = tf.keras.optimizers.Adam(learning_rate=0.0001) ## Default is 0.001
B1_model.compile(optimizer=opt, loss='categorical_crossentropy',
metrics=['accuracy'])
```

##### 1.4.2 The Two-Branch Model

In the two-branch model (B2), an additional branch was added, which obtains the same input as the first unaltered branch. The architecture of this second branch is the same as that of the first branch, except for the shapes of the convolutional filter kernels. In this branch, the first kernel dimension was set to 2, i.e. each kernel spans two rows. Note that we follow the (counterintuitive) TensorFlow notation, in which kernel shapes are specified in the format (height, width). The second dimension was chosen to be equal to the square-shaped kernels in the corresponding contraction and expansion paths. This leads to kernel shapes of 2×11, 2×7, 2×5,2×4, 2×3 in the contraction and kernel shapes of 2×4, 2×5, 2×7, 2×11 in the expansion path. The purpose is to train the kernels to keep the nucleotide order intact. Again the same code was used for the 48×48 as well as the 96×96 study cases. Output shapes in the following code depend on whether the input size is 48×48 or 96×96. To better understand the encoder and decoder path, we added the output shape of some layers as comments for the 48×48 study cases.

**Code of the B2 Model Architecture**

```
numRows  = 48        # OR 96
numCols  = 48        # OR 96
act_fun  = “sigmoid” # OR “relu”
act_init = “glorot_normal” # OR “he_normal”

inputs = tf.keras.layers.Input(shape=(numRows, numCols, 5))
#Contraction path
c1 = tf.keras.layers.Conv2D(32, (2, 11), activation=act_fun,
kernel_initializer=act_init, padding='same')(inputs)
# Output shape: 48x48x32

c1 = tf.keras.layers.Dropout(0.1)(c1)
c1 = tf.keras.layers.Conv2D(32, (2, 11), activation=act_fun,
kernel_initializer=act_init, padding='same')(c1)
p1 = tf.keras.layers.MaxPooling2D((2, 2))(c1)
# Output shape: 24x24x32

c2 = tf.keras.layers.Conv2D(64, (2, 7), activation=act_fun,
kernel_initializer=act_init, padding='same')(p1)
# Output shape: 24x24x64
c2 = tf.keras.layers.Dropout(0.1)(c2)
c2 = tf.keras.layers.Conv2D(64, (2, 7), activation=act_fun,
kernel_initializer=act_init, padding='same')(c2)
p2 = tf.keras.layers.MaxPooling2D((2, 2))(c2)
# Output shape: 12x12x64

c3 = tf.keras.layers.Conv2D(128, (2, 5), activation=act_fun,
kernel_initializer=act_init, padding='same')(p2)
# Output shape: 12x12x128
c3 = tf.keras.layers.Dropout(0.2)(c3)
c3 = tf.keras.layers.Conv2D(128, (2, 5), activation=act_fun,
```

```
kernel_initializer=act_init, padding='same')(c3)
p3 = tf.keras.layers.MaxPooling2D((2, 2))(c3)
# Output shape: 6x6x128

c4 = tf.keras.layers.Conv2D(128, (2, 4), activation=act_fun,
kernel_initializer=act_init, padding='same')(p3)
#Output shape: 6x6x128
c4 = tf.keras.layers.Dropout(0.2)(c4)
c4 = tf.keras.layers.Conv2D(128, (2, 4), activation=act_fun,
kernel_initializer=act_init, padding='same')(c4)
p4 = tf.keras.layers.MaxPooling2D(pool_size=(2, 2))(c4)
# Output shape: 3x3x128

c5 = tf.keras.layers.Conv2D(256, (2, 3), activation=act_fun,
kernel_initializer=act_init, padding='same')(p4)
c5 = tf.keras.layers.Dropout(0.3)(c5)
c5 = tf.keras.layers.Conv2D(256, (2, 3), activation=act_fun,
kernel_initializer=act_init, padding='same')(c5)
# Output shape: 3x3x256

#Expansive path
u6 = tf.keras.layers.Conv2DTranspose(128, (2, 2), strides=(2, 2),
padding='same')(c5)
# Output shape: 6x6x128
u6 = tf.keras.layers.concatenate([u6, c4])
# Output shape is 6x6x256
c6 = tf.keras.layers.Conv2D(128, (2, 4), activation=act_fun,
kernel_initializer=act_init, padding='same')(u6)
# Output shape is 6x6x128
c6 = tf.keras.layers.Dropout(0.2)(c6)
c6 = tf.keras.layers.Conv2D(128, (2, 4), activation=act_fun,
kernel_initializer=act_init, padding='same')(c6)
# Output shape is 6x6x128

u7 = tf.keras.layers.Conv2DTranspose(128, (2, 2), strides=(2, 2),
padding='same')(c6)
# Output shape: 12x12x128
u7 = tf.keras.layers.concatenate([u7, c3])
# Output shape: 12x12x256
c7 = tf.keras.layers.Conv2D(128, (2, 5), activation=act_fun,
kernel_initializer=act_init, padding='same')(u7)
c7 = tf.keras.layers.Dropout(0.2)(c7)
c7 = tf.keras.layers.Conv2D(128, (2, 5), activation=act_fun,
kernel_initializer=act_init, padding='same')(c7)
# Output shape: 12x12x128

u8 = tf.keras.layers.Conv2DTranspose(64, (2, 2), strides=(2, 2),
padding='same')(c7)
# Output shape: 24x24x64

u8 = tf.keras.layers.concatenate([u8, c2])
# Output shape: 24x24x128
c8 = tf.keras.layers.Conv2D(64, (2, 7), activation=act_fun,
kernel_initializer=act_init, padding='same')(u8)
c8 = tf.keras.layers.Dropout(0.1)(c8)
c8 = tf.keras.layers.Conv2D(64, (2, 7), activation=act_fun,
kernel_initializer=act_init, padding='same')(c8)
# Output shape: 24x24x64
```

```
u9 = tf.keras.layers.Conv2DTranspose(32, (2, 2), strides=(2, 2),
padding='same')(c8)
# Output shape: 48x48x32
u9 = tf.keras.layers.concatenate([u9, c1], axis=3)
# Output shape is 48x48x64
c9 = tf.keras.layers.Conv2D(32, (2, 11), activation=act_fun,
kernel_initializer=act_init, padding='same')(u9)
c9 = tf.keras.layers.Dropout(0.1)(c9)
c9 = tf.keras.layers.Conv2D(32, (2, 11), activation=act_fun,
kernel_initializer=act_init, padding='same')(c9)
# Output shape: 48x48x32
outputs = tf.keras.layers.Conv2D(5, (1, 1), activation='sigmoid')(c9)
y = tf.keras.Model(inputs=inputs,outputs=outputs)
# Output shape: 48x48x5

#Input 2
#Contraction path
c1 = tf.keras.layers.Conv2D(32, (11, 11), activation=act_fun,
kernel_initializer=act_init, padding='same')(inputs)
# Output shape: 48x48x32
c1 = tf.keras.layers.Dropout(0.1)(c1)
c1 = tf.keras.layers.Conv2D(32, (11, 11), activation=act_fun,
kernel_initializer=act_init, padding='same')(c1)
p1 = tf.keras.layers.MaxPooling2D((2, 2))(c1)
# Output shape: 24x24x32

c2 = tf.keras.layers.Conv2D(64, (7, 7), activation=act_fun,
kernel_initializer=act_init, padding='same')(p1)
# Output shape: 24x24x64
c2 = tf.keras.layers.Dropout(0.1)(c2)
c2 = tf.keras.layers.Conv2D(64, (7, 7), activation=act_fun,
kernel_initializer=act_init, padding='same')(c2)
p2 = tf.keras.layers.MaxPooling2D((2, 2))(c2)
# Output shape: 12x12x64

c3 = tf.keras.layers.Conv2D(128, (5, 5), activation=act_fun,
kernel_initializer=act_init, padding='same')(p2)
# Output shape: 12x12x128
c3 = tf.keras.layers.Dropout(0.2)(c3)
c3 = tf.keras.layers.Conv2D(128, (5, 5), activation=act_fun,
kernel_initializer=act_init, padding='same')(c3)
p3 = tf.keras.layers.MaxPooling2D((2, 2))(c3)
# Output shape: 6x6x128

c4 = tf.keras.layers.Conv2D(128, (4, 4), activation=act_fun,
kernel_initializer=act_init, padding='same')(p3)
# Output shape: 6x6x128
c4 = tf.keras.layers.Dropout(0.2)(c4)
c4 = tf.keras.layers.Conv2D(128, (4, 4), activation=act_fun,
kernel_initializer=act_init, padding='same')(c4)
p4 = tf.keras.layers.MaxPooling2D(pool_size=(2, 2))(c4)
# Output shape: 3x3x128

c5 = tf.keras.layers.Conv2D(256, (3, 3), activation=act_fun,
kernel_initializer=act_init, padding='same')(p4)
c5 = tf.keras.layers.Dropout(0.3)(c5)
c5 = tf.keras.layers.Conv2D(256, (3, 3), activation=act_fun,
kernel_initializer=act_init, padding='same')(c5)
# Output shape: 3x3x256
```

```
#Expansive path
u6 = tf.keras.layers.Conv2DTranspose(128, (2, 2), strides=(2, 2),
padding='same')(c5)
# Output shape: 6x6x128
u6 = tf.keras.layers.concatenate([u6, c4])
# Output shape: 6x6x256
c6 = tf.keras.layers.Conv2D(128, (4, 4), activation=act_fun,
kernel_initializer=act_init, padding='same')(u6)
c6 = tf.keras.layers.Dropout(0.2)(c6)
c6 = tf.keras.layers.Conv2D(128, (4, 4), activation=act_fun,
kernel_initializer=act_init, padding='same')(c6)
# Output shape: 6x6x128

u7 = tf.keras.layers.Conv2DTranspose(128, (2, 2), strides=(2, 2),
padding='same')(c6)
# Output shape: 12x12x128
u7 = tf.keras.layers.concatenate([u7, c3])
# Output shape: 12x12x256
c7 = tf.keras.layers.Conv2D(128, (5, 5), activation=act_fun,
kernel_initializer=act_init, padding='same')(u7)
c7 = tf.keras.layers.Dropout(0.2)(c7)
c7 = tf.keras.layers.Conv2D(128, (5, 5), activation=act_fun,
kernel_initializer=act_init, padding='same')(c7)
# Output shape: 12x12x128

u8 = tf.keras.layers.Conv2DTranspose(64, (2, 2), strides=(2, 2),
padding='same')(c7)
# Output shape: 24x24x64
u8 = tf.keras.layers.concatenate([u8, c2])
# Output shape: 24x24x128
c8 = tf.keras.layers.Conv2D(64, (7, 7), activation=act_fun,
kernel_initializer=act_init, padding='same')(u8)
c8 = tf.keras.layers.Dropout(0.1)(c8)
c8 = tf.keras.layers.Conv2D(64, (7, 7), activation=act_fun,
kernel_initializer=act_init, padding='same')(c8)
# Output shape: 24x24x64

u9 = tf.keras.layers.Conv2DTranspose(32, (2, 2), strides=(2, 2),
padding='same')(c8)
# Output shape: 48x48x32
u9 = tf.keras.layers.concatenate([u9, c1], axis=3)
# Output shape: 48x48x64
c9 = tf.keras.layers.Conv2D(32, (11, 11), activation=act_fun,
kernel_initializer=act_init, padding='same')(u9)
c9 = tf.keras.layers.Dropout(0.1)(c9)
c9 = tf.keras.layers.Conv2D(32, (11, 11), activation=act_fun,
kernel_initializer=act_init, padding='same')(c9)
# Output shape: 48x48x32
outputs = tf.keras.layers.Conv2D(5, (1, 1), activation='sigmoid')(c9)
z = tf.keras.Model(inputs=inputs,outputs=outputs)
# Output shape: 48x48x5

combined = tf.keras.layers.concatenate([y.output, z.output])
# Output shape: 48x48x10
a = tf.keras.layers.Conv2D(5, (1, 1), activation="softmax")(combined)
# Output shape: 48x48x5

B2_model = tf.keras.Model(inputs=[inputs],outputs=a)
```

```
# Output shape: 48x48x5

# Create and compile the model# Adam is chosen as an optimizer with a
learning rate of 0.0001

opt = tf.keras.optimizers.Adam(learning_rate=0.0001) ## Default is 0.001
B2_model.compile(optimizer=opt, loss='categorical_crossentropy',
metrics=['accuracy'])
```

##### 1.4.3 Three-Branch Model

Finally, in the three-branch model (B3) a third branch is added in which each filter kernel spans two columns and the same number of rows as the corresponding filters of the first branch with square-shaped kernels. Due to the increased size of the concatenated output of the three branches, two more convolutional layers are added to downsample the concatenated feature maps to five output channels.

###### Code of the B3 Model Architecture

```
numRows  = 48        # OR 96
numCols  = 48        # OR 96
act_fun  = "sigmoid" # OR "relu"
act_init = "glorot_normal" # OR "he_normal"

#Let's adapt the net to our model
inputs = tf.keras.layers.Input(shape=(args.rows, args.columns, 5))
#Input 1
#Contraction path
c1 = tf.keras.layers.Conv2D(32, (11, 2), activation=act_fun,
kernel_initializer=act_init, padding='same')(inputs)
# Output shape: 48x48x32
c1 = tf.keras.layers.Dropout(0.1)(c1)
c1 = tf.keras.layers.Conv2D(32, (11, 2), activation=act_fun,
kernel_initializer=act_init, padding='same')(c1)
p1 = tf.keras.layers.MaxPooling2D((2, 2))(c1)
# Output shape: 24x24x32

c2 = tf.keras.layers.Conv2D(64, (7, 2), activation=act_fun,
kernel_initializer=act_init, padding='same')(p1)
# Output shape: 24x24x64
c2 = tf.keras.layers.Dropout(0.1)(c2)
c2 = tf.keras.layers.Conv2D(64, (7, 2), activation=act_fun,
kernel_initializer=act_init, padding='same')(c2)
p2 = tf.keras.layers.MaxPooling2D((2, 2))(c2)
# Output shape: 12x12x64

c3 = tf.keras.layers.Conv2D(128, (5, 2), activation=act_fun,
kernel_initializer=act_init, padding='same')(p2)
# Output shape: 12x12x128
c3 = tf.keras.layers.Dropout(0.2)(c3)
c3 = tf.keras.layers.Conv2D(128, (5, 2), activation=act_fun,
kernel_initializer=act_init, padding='same')(c3)
p3 = tf.keras.layers.MaxPooling2D((2, 2))(c3)
# Output shape: 6x6x128
```

```
c4 = tf.keras.layers.Conv2D(128, (4, 2), activation=act_fun,
kernel_initializer=act_init, padding='same')(p3)
# Output shape: 6x6x128
c4 = tf.keras.layers.Dropout(0.2)(c4)
c4 = tf.keras.layers.Conv2D(128, (4, 2), activation=act_fun,
kernel_initializer=act_init, padding='same')(c4)
p4 = tf.keras.layers.MaxPooling2D(pool_size=(2, 2))(c4)
# Output shape: 3x3x128

c5 = tf.keras.layers.Conv2D(256, (3, 2), activation=act_fun,
kernel_initializer=act_init, padding='same')(p4)
# Output shape: 3x3x256
c5 = tf.keras.layers.Dropout(0.3)(c5)
c5 = tf.keras.layers.Conv2D(256, (3, 2), activation=act_fun,
kernel_initializer=act_init, padding='same')(c5)
# Output shape: 3x3x256

#Expansive path
u6 = tf.keras.layers.Conv2DTranspose(128, (2, 2), strides=(2, 2),
padding='same')(c5)
# Output shape: 6x6x128
u6 = tf.keras.layers.concatenate([u6, c4])
# Output shape: 6x6x256
c6 = tf.keras.layers.Conv2D(128, (4, 2), activation=act_fun,
kernel_initializer=act_init, padding='same')(u6)
c6 = tf.keras.layers.Dropout(0.2)(c6)
c6 = tf.keras.layers.Conv2D(128, (4, 2), activation=act_fun,
kernel_initializer=act_init, padding='same')(c6)
# Output shape: 6x6x128

u7 = tf.keras.layers.Conv2DTranspose(128, (2, 2), strides=(2, 2),
padding='same')(c6)
# Output shape: 12x12x128
u7 = tf.keras.layers.concatenate([u7, c3])
# Output shape: 12x12x256
c7 = tf.keras.layers.Conv2D(128, (5, 2), activation=act_fun,
kernel_initializer=act_init, padding='same')(u7)
c7 = tf.keras.layers.Dropout(0.2)(c7)
c7 = tf.keras.layers.Conv2D(128, (5, 2), activation=act_fun,
kernel_initializer=act_init, padding='same')(c7)
# Output shape: 12x12x128

u8 = tf.keras.layers.Conv2DTranspose(64, (2, 2), strides=(2, 2),
padding='same')(c7)
# Output shape: 24x24x64
u8 = tf.keras.layers.concatenate([u8, c2])
# Output shape: 24x24x128
c8 = tf.keras.layers.Conv2D(64, (7, 2), activation=act_fun,
kernel_initializer=act_init, padding='same')(u8)
c8 = tf.keras.layers.Dropout(0.1)(c8)
c8 = tf.keras.layers.Conv2D(64, (7, 2), activation=act_fun,
kernel_initializer=act_init, padding='same')(c8)
# Output shape: 24x24x64

u9 = tf.keras.layers.Conv2DTranspose(32, (2, 2), strides=(2, 2),
padding='same')(c8)
# Output shape: 48x48x32
u9 = tf.keras.layers.concatenate([u9, c1], axis=3)
# Output shape: 48x48x64
```

```
c9 = tf.keras.layers.Conv2D(32, (11, 2), activation=act_fun,
kernel_initializer=act_init, padding='same')(u9)
c9 = tf.keras.layers.Dropout(0.1)(c9)
c9 = tf.keras.layers.Conv2D(32, (11, 2), activation=act_fun,
kernel_initializer=act_init, padding='same')(c9)
# Output shape: 48x48x32
outputs = tf.keras.layers.Conv2D(5, (1, 1), activation='sigmoid')(c9)
x = tf.keras.Model(inputs=inputs,outputs=outputs)
# Output shape: 48x48x5

#Input 2:
# Contraction path
c1 = tf.keras.layers.Conv2D(32, (2, 11), activation=act_fun,
kernel_initializer=act_init, padding='same')(inputs)
# Output shape: 48x48x32
c1 = tf.keras.layers.Dropout(0.1)(c1)
c1 = tf.keras.layers.Conv2D(32, (2, 11), activation=act_fun,
kernel_initializer=act_init, padding='same')(c1)
p1 = tf.keras.layers.MaxPooling2D((2, 2))(c1)
# Output shape: 24x24x32

c2 = tf.keras.layers.Conv2D(64, (2, 7), activation=act_fun,
kernel_initializer=act_init, padding='same')(p1)
# Output shape: 24x24x64
c2 = tf.keras.layers.Dropout(0.1)(c2)
c2 = tf.keras.layers.Conv2D(64, (2, 7), activation=act_fun,
kernel_initializer=act_init, padding='same')(c2)
p2 = tf.keras.layers.MaxPooling2D((2, 2))(c2)
# Output shape: 12x12x64

c3 = tf.keras.layers.Conv2D(128, (2, 5), activation=act_fun,
kernel_initializer=act_init, padding='same')(p2)
# Output shape: 12x12x128
c3 = tf.keras.layers.Dropout(0.2)(c3)
c3 = tf.keras.layers.Conv2D(128, (2, 5), activation=act_fun,
kernel_initializer=act_init, padding='same')(c3)
p3 = tf.keras.layers.MaxPooling2D((2, 2))(c3)
# Output shape: 6x6x128

c4 = tf.keras.layers.Conv2D(128, (2, 4), activation=act_fun,
kernel_initializer=act_init, padding='same')(p3)
# Output shape: 6x6x128
c4 = tf.keras.layers.Dropout(0.2)(c4)
c4 = tf.keras.layers.Conv2D(128, (2, 4), activation=act_fun,
kernel_initializer=act_init, padding='same')(c4)
p4 = tf.keras.layers.MaxPooling2D(pool_size=(2, 2))(c4)
# Output shape: 3x3x128

c5 = tf.keras.layers.Conv2D(256, (2, 3), activation=act_fun,
kernel_initializer=act_init, padding='same')(p4)
# Output shape: 3x3x256
c5 = tf.keras.layers.Dropout(0.3)(c5)
c5 = tf.keras.layers.Conv2D(256, (2, 3), activation=act_fun,
kernel_initializer=act_init, padding='same')(c5)
# Output shape: 3x3x256

#Expansive path
u6 = tf.keras.layers.Conv2DTranspose(128, (2, 2), strides=(2, 2),
padding='same')(c5)
```

```
# Output shape: 6x6x128
u6 = tf.keras.layers.concatenate([u6, c4])
# Output shape: 6x6x256
c6 = tf.keras.layers.Conv2D(128, (2, 4), activation=act_fun,
kernel_initializer=act_init, padding='same')(u6)
c6 = tf.keras.layers.Dropout(0.2)(c6)
c6 = tf.keras.layers.Conv2D(128, (2, 4), activation=act_fun,
kernel_initializer=act_init, padding='same')(c6)
# Output shape: 6x6x128

u7 = tf.keras.layers.Conv2DTranspose(128, (2, 2), strides=(2, 2),
padding='same')(c6)
# Output shape: 12x12x128
u7 = tf.keras.layers.concatenate([u7, c3])
# Output shape: 12x12x256
c7 = tf.keras.layers.Conv2D(128, (2, 5), activation=act_fun,
kernel_initializer=act_init, padding='same')(u7)
c7 = tf.keras.layers.Dropout(0.2)(c7)
c7 = tf.keras.layers.Conv2D(128, (2, 5), activation=act_fun,
kernel_initializer=act_init, padding='same')(c7)
# Output shape: 12x12x128

u8 = tf.keras.layers.Conv2DTranspose(64, (2, 2), strides=(2, 2),
padding='same')(c7)
# Output shape: 24x24x64
u8 = tf.keras.layers.concatenate([u8, c2])
# Output shape: 24x24x128
c8 = tf.keras.layers.Conv2D(64, (2, 7), activation=act_fun,
kernel_initializer=act_init, padding='same')(u8)
c8 = tf.keras.layers.Dropout(0.1)(c8)
c8 = tf.keras.layers.Conv2D(64, (2, 7), activation=act_fun,
kernel_initializer=act_init, padding='same')(c8)
# Output shape: 24x24x64

u9 = tf.keras.layers.Conv2DTranspose(32, (2, 2), strides=(2, 2),
padding='same')(c8)
# Output shape: 48x48x32
u9 = tf.keras.layers.concatenate([u9, c1], axis=3)
# Output shape: 48x48x64
c9 = tf.keras.layers.Conv2D(32, (2, 11), activation=act_fun,
kernel_initializer=act_init, padding='same')(u9)
c9 = tf.keras.layers.Dropout(0.1)(c9)
c9 = tf.keras.layers.Conv2D(32, (2, 11), activation=act_fun,
kernel_initializer=act_init, padding='same')(c9)
# Output shape: 48x48x32
outputs = tf.keras.layers.Conv2D(5, (1, 1), activation='sigmoid')(c9)
y = tf.keras.Model(inputs=inputs,outputs=outputs)
# Output shape: 48x48x5

#Input 3
#Contraction path
c1 = tf.keras.layers.Conv2D(32, (11, 11), activation=act_fun,
kernel_initializer=act_init, padding='same')(inputs)
# Output shape: 48x48x32
c1 = tf.keras.layers.Dropout(0.1)(c1)
c1 = tf.keras.layers.Conv2D(32, (11, 11), activation=act_fun,
kernel_initializer=act_init, padding='same')(c1)
p1 = tf.keras.layers.MaxPooling2D((2, 2))(c1)
# Output shape: 24x24x32
```

```
c2 = tf.keras.layers.Conv2D(64, (7, 7), activation=act_fun,
kernel_initializer=act_init, padding='same')(p1)
# Output shape: 24x24x64
c2 = tf.keras.layers.Dropout(0.1)(c2)
c2 = tf.keras.layers.Conv2D(64, (7, 7), activation=act_fun,
kernel_initializer=act_init, padding='same')(c2)
p2 = tf.keras.layers.MaxPooling2D((2, 2))(c2)
# Output shape: 12x12x64

c3 = tf.keras.layers.Conv2D(128, (5, 5), activation=act_fun,
kernel_initializer=act_init, padding='same')(p2)
# Output shape: 12x12x128
c3 = tf.keras.layers.Dropout(0.2)(c3)
c3 = tf.keras.layers.Conv2D(128, (5, 5), activation=act_fun,
kernel_initializer=act_init, padding='same')(c3)
p3 = tf.keras.layers.MaxPooling2D((2, 2))(c3)
# Output shape: 6x6x128

c4 = tf.keras.layers.Conv2D(128, (4, 4), activation=act_fun,
kernel_initializer=act_init, padding='same')(p3)
# Output shape: 6x6x128
c4 = tf.keras.layers.Dropout(0.2)(c4)
c4 = tf.keras.layers.Conv2D(128, (4, 4), activation=act_fun,
kernel_initializer=act_init, padding='same')(c4)
p4 = tf.keras.layers.MaxPooling2D(pool_size=(2, 2))(c4)
# Output shape: 3x3x128

c5 = tf.keras.layers.Conv2D(256, (3, 3), activation=act_fun,
kernel_initializer=act_init, padding='same')(p4)
# Output shape: 3x3x256
c5 = tf.keras.layers.Dropout(0.3)(c5)
c5 = tf.keras.layers.Conv2D(256, (3, 3), activation=act_fun,
kernel_initializer=act_init, padding='same')(c5)
# Output shape: 3x3x256

#Expansive path
u6 = tf.keras.layers.Conv2DTranspose(128, (2, 2), strides=(2, 2),
padding='same')(c5)
# Output shape: 6x6x128
u6 = tf.keras.layers.concatenate([u6, c4])
# Output shape: 6x6x256
c6 = tf.keras.layers.Conv2D(128, (4, 4), activation=act_fun,
kernel_initializer=act_init, padding='same')(u6)
c6 = tf.keras.layers.Dropout(0.2)(c6)
c6 = tf.keras.layers.Conv2D(128, (4, 4), activation=act_fun,
kernel_initializer=act_init, padding='same')(c6)
# Output shape: 6x6x128

u7 = tf.keras.layers.Conv2DTranspose(128, (2, 2), strides=(2, 2),
padding='same')(c6)
# Output shape: 12x12x128
u7 = tf.keras.layers.concatenate([u7, c3])
# Output shape: 12x12x256
c7 = tf.keras.layers.Conv2D(128, (5, 5), activation=act_fun,
kernel_initializer=act_init, padding='same')(u7)
c7 = tf.keras.layers.Dropout(0.2)(c7)
c7 = tf.keras.layers.Conv2D(128, (5, 5), activation=act_fun,
kernel_initializer=act_init, padding='same')(c7)
```

```
# Output shape: 12x12x128

u8 = tf.keras.layers.Conv2DTranspose(64, (2, 2), strides=(2, 2),
padding='same')(c7)
# Output shape: 24x24x64
u8 = tf.keras.layers.concatenate([u8, c2])
# Output shape: 24x24x128
c8 = tf.keras.layers.Conv2D(64, (7, 7), activation=act_fun,
kernel_initializer=act_init, padding='same')(u8)
c8 = tf.keras.layers.Dropout(0.1)(c8)
c8 = tf.keras.layers.Conv2D(64, (7, 7), activation=act_fun,
kernel_initializer=act_init, padding='same')(c8)
# Output shape: 24x24x64

u9 = tf.keras.layers.Conv2DTranspose(32, (2, 2), strides=(2, 2),
padding='same')(c8)
# Output shape: 48x48x32
u9 = tf.keras.layers.concatenate([u9, c1], axis=3)
# Output shape: 48x48x64
c9 = tf.keras.layers.Conv2D(32, (11, 11), activation=act_fun,
kernel_initializer=act_init, padding='same')(u9)
c9 = tf.keras.layers.Dropout(0.1)(c9)
c9 = tf.keras.layers.Conv2D(32, (11, 11), activation=act_fun,
kernel_initializer=act_init, padding='same')(c9)
# Output shape: 48x48x32
outputs = tf.keras.layers.Conv2D(5, (1, 1), activation='sigmoid')(c9)
z = tf.keras.Model(inputs=inputs,outputs=outputs)
# Output shape: 48x48x5

combined = tf.keras.layers.concatenate([x.output, y.output, z.output])
# Output shape: 48x48x15
a = tf.keras.layers.Conv2D(10, (1, 1), activation="relu")(combined)
a = tf.keras.layers.Conv2D(5, (1, 1), activation="softmax")(a)"
# Output shape: 48x48x5

B3_model = tf.keras.Model(inputs=[inputs],outputs=a)
# Output shape: 48x48x5

# Create and compile the model# Adam is chosen as an optimizer with a
learning rate of 0.0001

opt = tf.keras.optimizers.Adam(learning_rate=0.0001) ## Default is 0.001
B3_model.compile(optimizer=opt, loss='categorical_crossentropy',
metrics=['accuracy'])
```

#### 1.5 Numerical stability and kernel initialization

In the present project, we used and compared the sigmoid and ReLu activation functions. Depending on the activation function, we sometimes needed multiple trials until the neural network started to learn. We were able to avoid this numerical problem by choosing the activation function-specific kernel initializers `he_normal` for ReLU and `glorot_normal` for sigmoid.

##### 1.6 Technique used to combine any number of feature maps to five output channels

The standard technique to combine any number of feature maps to smaller values, such as three output channels in the case of images or five output channels in the present study, is to use a two-dimensional convolutional layer with kernel size 1×1 and N filter kernels, where N is the number of features the output should be reduced to. That is N=3 for images and N=5 for five different output dimensions as in our case. In Tensorflow we use the `Conv2D(5,(1,1))` layer. This layer has $F_{out}\times(F_{in}+1)$ trainable parameters over the whole matrix, where $F_{in}$ is the number of feature maps that have to be combined and $F_{out}=3$ or $F_{out}=5$ is the number of output channels. In the case of the B1, B2, and B3 models we used this approach to combine the feature maps of each branch into 5 output channels. Furthermore, we used this to combine the concatenated feature maps in the B2 or B3 model and reduce this to five output channels.

### 2 Supplementary Results

#### 2.1 Visualizing the Output

Alignment performance and complexity of the alignment problem are best visualized by comparing unaligned sequences with aligned sequences. To give the reader an impression of what kind of alignments have been tested, we show in Figure S6 an example of a sequence matrix of unaligned nucleotide sequences and in Figure S7 the result after predicting the alignment for these sequences with the Ali-U-Net B3 model trained for the 48x48 RSE+IG study case. The example was chosen from the test dataset of this study case.

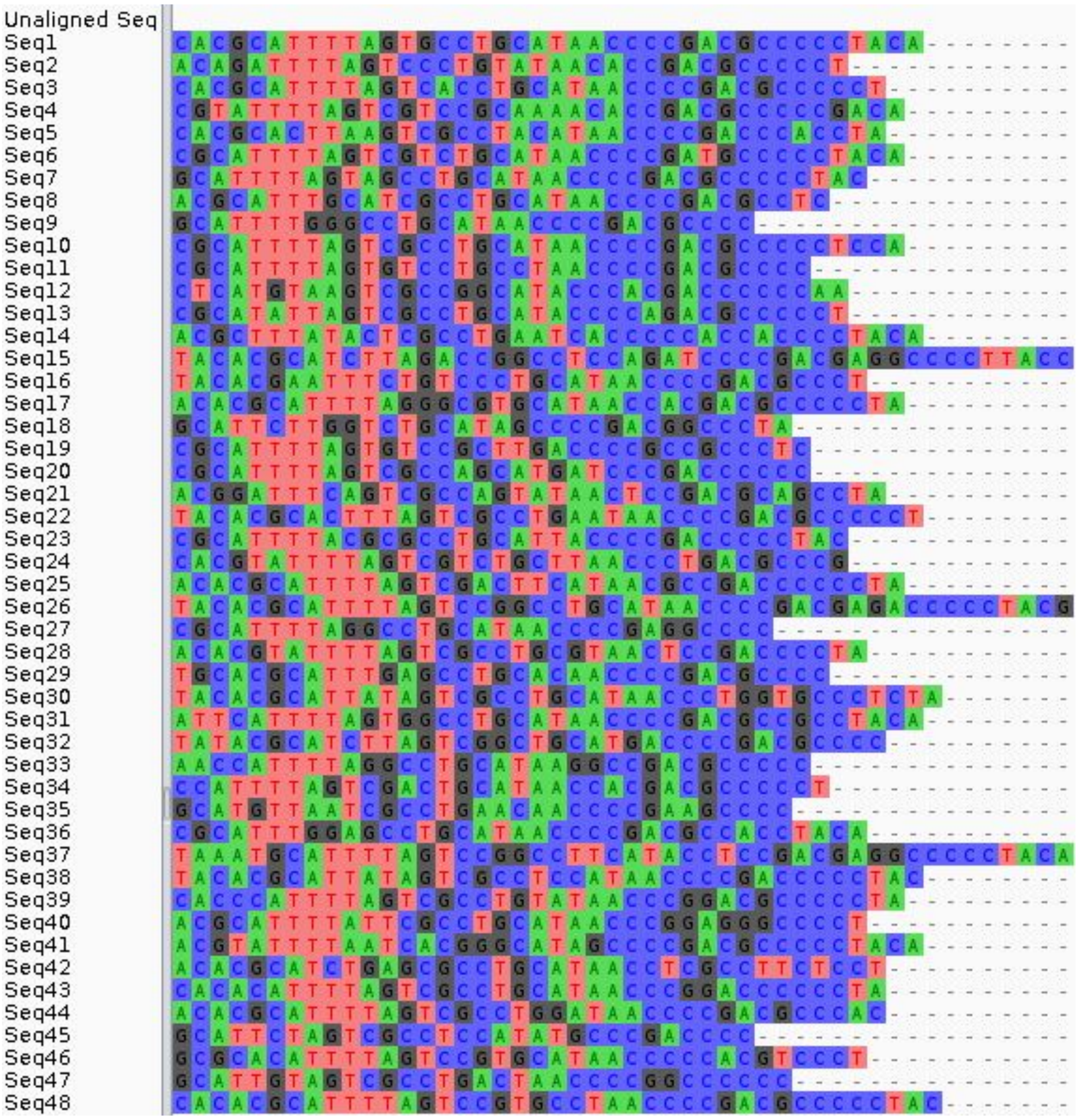

**Figure S6:** Unaligned reference sequences serving as input for the Ai-U-Net. The predicted alignment is shown in Figure S7.

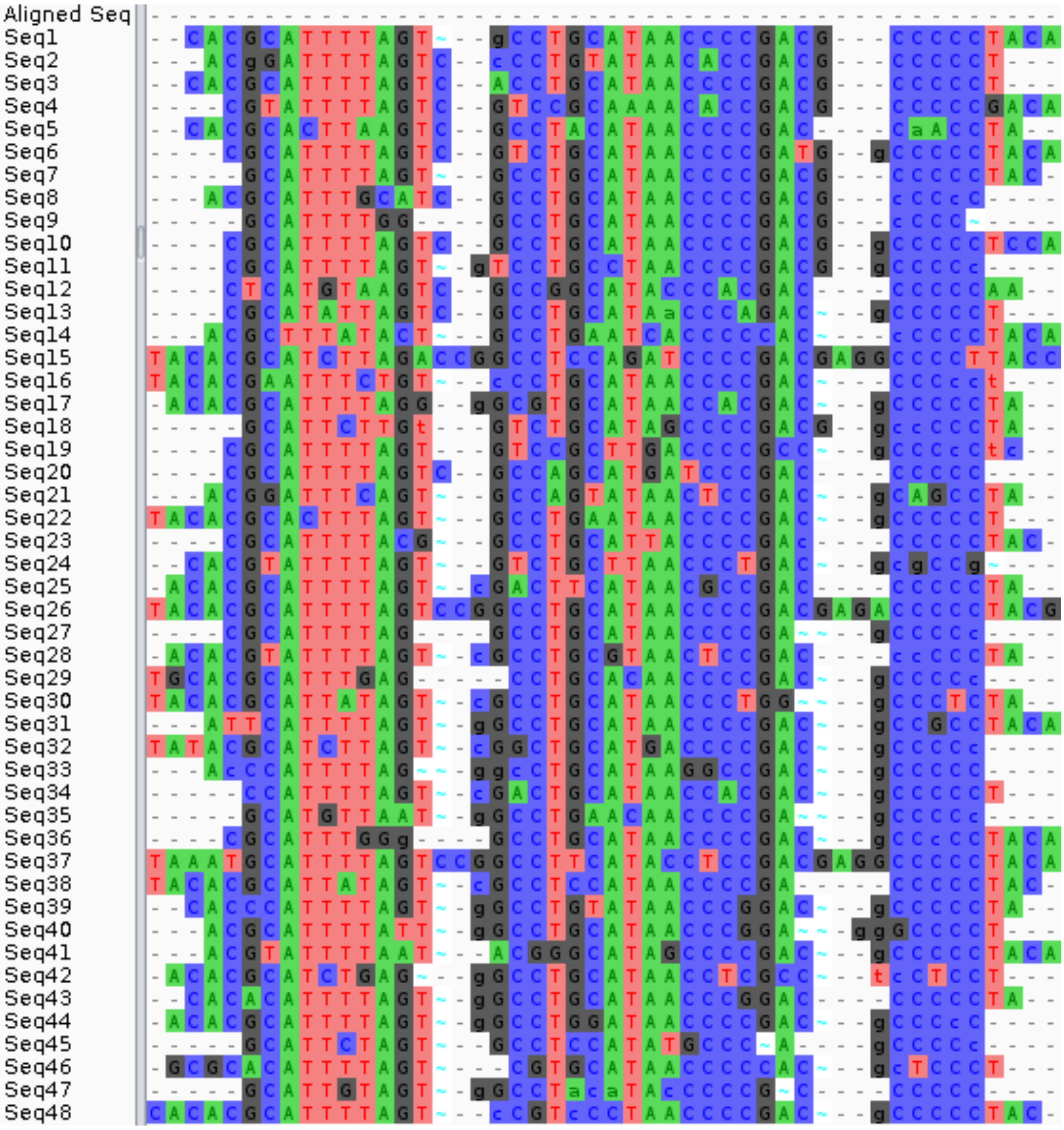


**Figure S7:** Example of aligned sequences predicted with the Ali-U-Net. Upper case letters and normal gap characters depict nucleotides and gaps that are identical to those in the reference alignment, while lower case and tilde characters depict differences. Differences can be due to alternative alignments or wrongly inserted bases or gaps, i.e. hallucinations.

#### 2.2 Validation of models trained for mixed study cases on unstructured gaps

We validated the trained neural networks on the test datasets that correspond to the training data sets for the study cases (Table 2). Furthermore, two Ali-U-Net$_{mix}$ models, one for 48x48 and one for 96x96 sequence matrices, have been trained on a mixture of RSE and RES+IG alignments and validated on the test datasets of the RES and RSE+IG study cases (Table 6). Now we evaluate the performance of the Ali-U-Net$_{mix}$ models on sequence matrices for which 1-3 unstructured gaps have been simulated. These unstructured gaps can occur with uniform probability across the sequence matrix and can span up to 20% of the sequence length. They can occur in regions of the nucleotide matrices in which the training data did not contain any gaps or only gaps with a block structure. Our test dataset consists of 1000 pairs of simulated reference alignments and the corresponding unaligned sequences. Our results show that the Ali-U-Net$_{mix}$ model does not align the sequences containing unstructured gaps properly. The problem is illustrated in Figure S8, which shows the total number of gap characters occurring at different alignment sites of the reference alignments and of the alignments predicted with the Ali-U-Net$_{mix}$ model for the case of 48x48 nucleotide matrices. The distributions show that gaps are systematically moved to the ends of the sequence since this is how the Ali-U-Net$_{mix}$ model was trained to deal with gaps that do not have a block-like structure. How a typical alignment looks in which the gaps are pushed to the end of the sequence can be seen in Figures S9 and S10. Figure S9 shows a simulated reference alignment with a nucleotide matrix of size 48x48. Figure S10 shows the alignment that the Ali-U-Net$_{mix}$ predicted when using the unaligned sequences of Figure S9 as input. Both gaps are pushed to the end of the sequences. The wrongly aligned nucleotides in the region between the correct gap positions and the position where the gaps have been inserted show an increased number of hallucinations, i.e. alterations of the original sequences.

For the unstructed gap test dataset, the overall alignment accuracy of the Ali-U-Net$_{mix}$ and that of the other widely used alignment programs is shown in Table S1. The overall mean accuracy of the Ali-U-Net$_{mix}$ model is 95.5% and 98.5% for the 48x48 and the 96x96 study cases. The high accuracy stems from the small number of instructed gaps in these alignments. Interestingly, the overall accuracy/identity of reference and predicted alignments is still higher for the Ali-U-Net$_{mix}$ models than for the MUSCLE software.

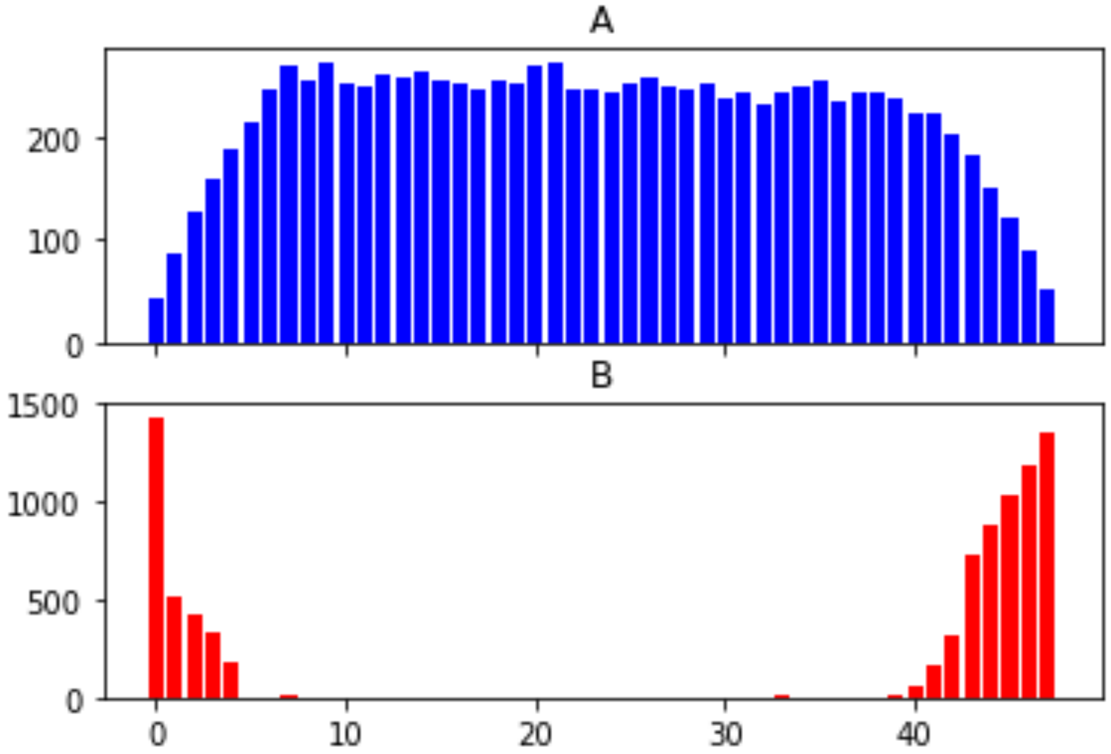


**Figure S8**: Total number of gap characters occurring at different alignment sites of (A) simulated reference alignments in the test dataset with unstructured gaps and (B) in the output alignments after predicting alignments with the Ali-U-Net$_{mix}$ model using the corresponding unaligned sequences in this test dataset as input. The results show that gaps are only inserted in columns in which the NN was trained to insert gaps.

| Model | 48x48 unstructured gaps test dataset | | 96x96 unstructured gaps test dataset | |
|---|---|---|---|---|
| | M | Mdn | M | Mdn |
| **Ali-U-Net B3$_{mix}$** trained with mixed data | 98.46 | 98.14 | 95.50 | 93.90 |
| **MAFFT** | 99.74 | 99.14 | 99.96 | 99.60 |
| **T-Coffee** | 99.89 | 99.91 | 99.90 | 99.98 |
| **Clustal-Omega** | 98.52 | 82.70 | 99.39 | 81.81 |
| **MUSCLE** | 80.99 | 81.08 | 81.14 | 81.17 |

**Table S1**: Comparison of the total identity scores of the Ali-U-Net$_{mix}$. (B3+ReLU), and four other commonly used sequence alignment programs when applied to two test datasets of thousand nucleotide matrices of size 96x96 and 48x48, simulated with unstructured gaps.

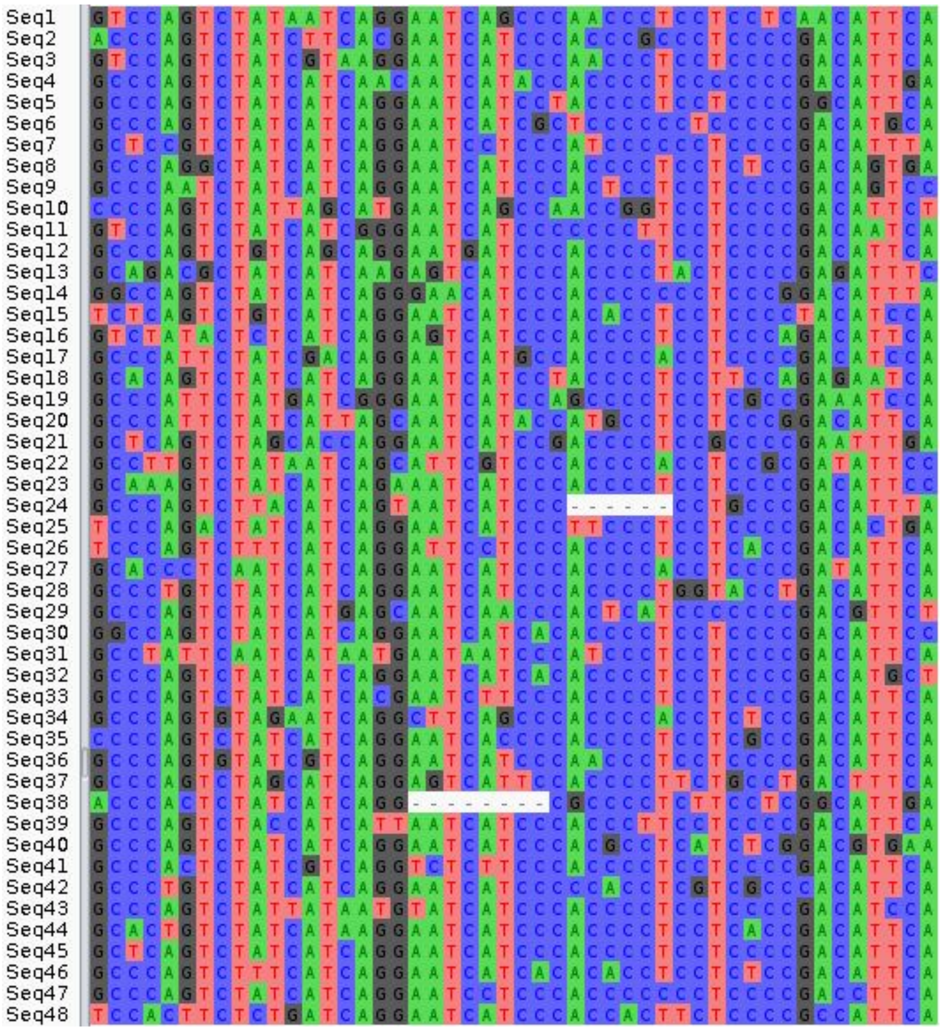


**Figure S9**: Example showing a typical simulated reference alignment from the test dataset with structured gaps, i.e. gaps that are allowed to occur uniformly across the alignment without block structure in a 48x48 window.

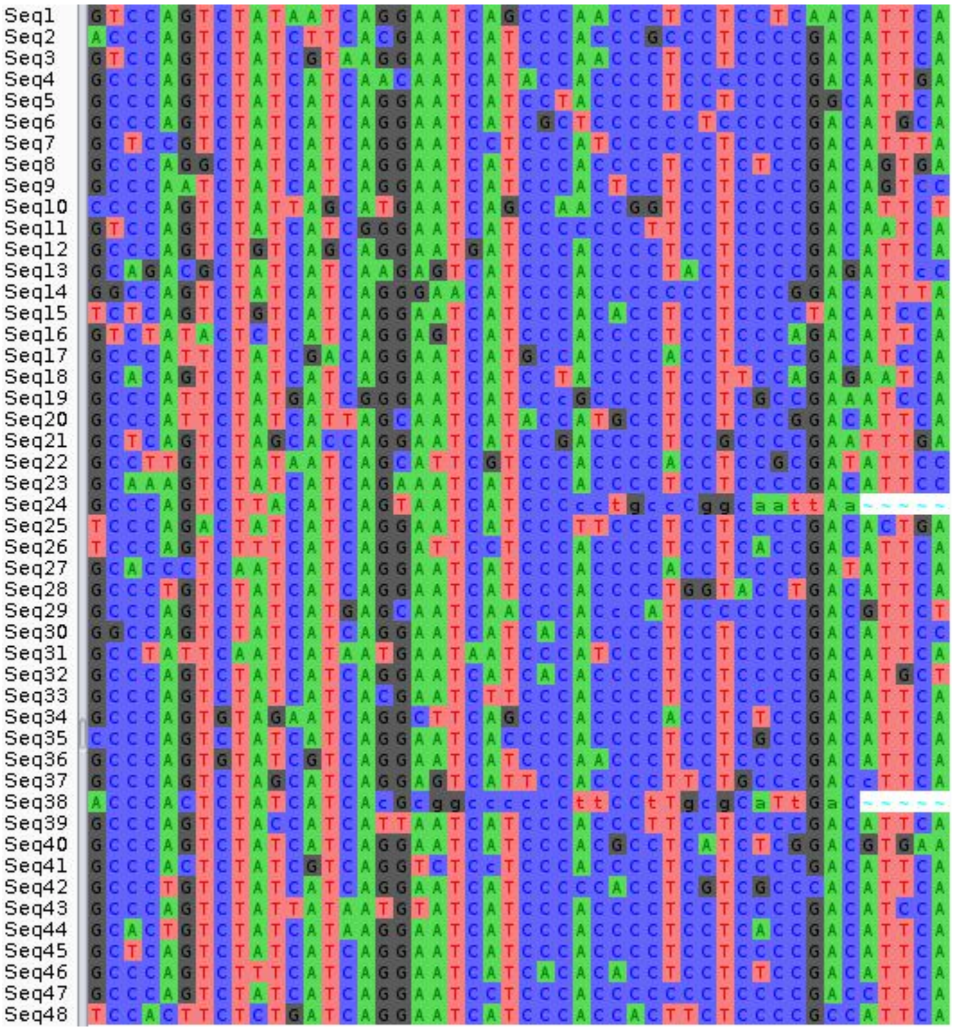


**Figure S10**: Alignment obtained with the Ali-U-Net$_{mix}$ model when using the unaligned sequences corresponding to the sequences in Figure S9 as input.
